# Molecular basis of unidirectional information transmission in two-component systems: lessons from the DesK-DesR thermosensor

**DOI:** 10.1101/2021.11.12.468217

**Authors:** Sofía Lima, Juan Blanco, Federico Olivieri, Juan Andrés Imelio, Federico Carrión, Beatriz Alvarez, Alejandro Buschiazzo, Marcelo Martí, Felipe Trajtenberg

## Abstract

Cellular signaling systems transmit information over long distances using allosteric transitions and/or post-translational modifications. In two-component systems the sensor histidine kinase and response regulator are wired through phosphoryl-transfer reactions, using either a uni- or bi-directional transmission mode, allowing to build rich regulatory networks. Using the thermosensor DesK-DesR two-component system from *Bacillus subtilis* and combining crystal structures, QM/MM calculations and integrative kinetic modeling, we uncover that: i) longer or shorter distances between the phosphoryl-acceptor and -donor residues can shift the phosphoryl-transfer equilibrium; ii) the phosphorylation-dependent dimerization of the regulator acts as a sequestering mechanism by preventing the interaction with the histidine kinase; and iii) the kinase’s intrinsic conformational equilibrium makes the phosphotransferase state unlikely in the absence of histidine phosphorylation, minimizing backwards transmission. These mechanisms allow the system to control the direction of signal transmission in a very efficient way, showcasing the key role that structure-encoded allostery plays in signaling proteins to store and transmit information.

## Introduction

Perception is a fundamental process that allows living organisms to adapt to highly variable environments. Cells are equipped with a set of signaling pathways that recognize specific signals, transmit and process the information to regulate different cellular programs. Two-component systems (TCS) are relevant constituents of this sensorial system in bacteria and archaea, also found in fungi and plants^1^. TCSs are usually composed of a sensor histidine kinase (HK), which detects a specific signal that allosterically regulates their catalytic activities. Information is then transmitted further downstream via phosphorylation of a specific response regulator (RR)^2^, which acts as the second component of the TCS pathway, executing the output response.

Upon the signal-dependent pathway activation, the HK autophosphorylates a conserved His. A second reaction takes place in tandem, and the phosphoryl moiety gets transferred to an invariant Asp within its cognate RR. Activation of the RR by phosphorylaton usually promotes its dimerization and subsequent binding to DNA, as most RRs bear a DNA-binding domain responsible for exerting the effector response by changing the expression of target genes^3^. In most TCSs, when the signal is absent, the HK also promotes the dephosphorylation of the RR (*i.e*. acting as a phoshatase), making of most HKs fascinating examples of paradoxical enzymes^4^. HK-mediated P~RR dephosphorylation has been shown to be an extremely relevant process for shutting down the system^5^ and preventing crosstalk^6^ *in vivo*. Interestingly, TCS pathways have also evolved to increase their complexity by wiring additional intermediate RR and His containing phosphotransferase proteins (HPt) domains that ultimately build phosphorylation cascades, known as phosphorelays.

In the past few years several groups have made important contributions to the understanding of the regulation of the activities of HK and RR^3,7-14^, how their components are functionally wired through specific protein-protein interactions^15^ and how the signal is transmitted from the extra- to the intra-cellular space^16,17^. Presently, five HK families (HisKA, HisKA_3, HisKA_2, H-kinase_dim and HWE_HK) have been recognized based on sequence clustering^18^; these families share many similarities at the structural level^9,11,13,19-21^. However, we are still far from understanding several fundamental aspects of how HK-driven phosphorylation cascades transmit and process information efficiently. For example, are phosphorylation/dephosphorylation futile cycles minimized to avoid cell energy dissipation? If so, how is this accomplished from a structural and mechanistic perspective? Or yet, how is the directionality of phosphoryl-transfer reactions enforced, such that the P~RR species are accumulated, or even to allow the connection with additional intermediate components to build richer phosphorelays pathways?

Phosphoryl-transfer reactions occur by nucleophilic substitution. The electron-rich oxygen of the RR’s Asp carboxylate, attacks the phosphorus electrophile, substituting the *ε*-nitrogen of the HK’s His imidazole^22^ (Extended Data Fig 1). The directionality issue should not be overlooked, since His phosphorylation is expected to be favored over Asp phosphorylation considering available data on the hydrolysis of phosphoramidate and phosphoanhydride compounds^23-25^. Dedicated mechanisms are thus anticipated to surmount the uphill direction and ensure proper physiologic behavior.

The TCS DesK-DesR from *Bacillus subtilis* is a thermosensor pathway that provides the cells with a fast homeostatic response that maintains cell membrane fluidity^26^. DesK is a HisKA_3 HK that is activated upon cold shock. Phosphorylation at the invariant D54_RR_ (Asp at position 54) of its cognate RR DesR triggers the adaptive response by activating the expression of a fatty acid desaturase^27^. The structural information available for this system is unique. Several experimental structures of both components in different steps of the signaling pathway, and adopting different conformations have been elucidated^7,9,28,29^. The catalytically active cytosolic and soluble region of DesK (DesKC) can adopt at least three functionally and structurally different conformations: autokinase, phosphatase and phosphotransferase states, involved in the autophosphorylation, phosphoryl-transfer and dephosphorylation activities, respectively. Based on the crystal structures of the DesK:DesR complex, and their comparison to other TCSs, the distance between the phosphoryl-donor and -acceptor residues (the HK His and RR’s Asp, respectively) correlates with the systems’ directionality or degree of phosphoryl-transfer reversibility. Shorter distances predict a tight transitions states in the nucleophilic substitution reaction^22^, *i.e*. a pentavalent bipyramidal transition state, and for still not understood reasons correlates with more reversible phosphoryl-transfer reactions. Inversely, longer distances between reactive residues imply loose transitions states, *i.e*. a more dissociated planar trigonal metaphosphate transition state, that correlate with irreversible P~His→Asp transfer reactions^7^.

In the present work, to further understand how phosphoryl-transfer reversibility is controlled in TCSs, quantum mechanics/molecular mechanics (QM/MM) calculations were performed describing the phosphoryl-transfer (phosphoryl transfer from His to Asp or Asp to His) and dephosphorylation reactions (phosphoryl-hydrolysis or transfer reaction from Asp to a water molecule) using the DesK-DesR system. The crystal structure of the DesK:DesR complex in the phosphatase state was refined to higher resolution than that previously available, further supporting the QM/MM data. Critical residues for the phosphotransferase and the phosphatase activities were pinpointed utilizing structure-guided point mutants. Finally, an integrative structural and kinetic model of the system was constructed, identifying key directionality determinants of the phosphoryl-transfer reaction. Overall, evidence that the different HK enzymatic activities are insulated from each other is provided, a pivotal element that ensures information is efficiently transmitted in the right direction.

## Results

### A hydroxyl anion is required for phospho-aspartate hydrolysis in the HK-mediated phosphatase reaction

To explore the mechanistic details of the dephosphorylation reaction in TCSs, the Free Energy Profile of the DesK-DesR system was calculated using QM/MM-based steered molecular dynamics. Calculations were based on an improved version of the crystal structure of the DesK:DesR complex in the phosphatase state (PDB id: 7SSJ), used as the starting species. The refinement of the previously reported structure was improved by reprocessing the raw diffraction data^30^ with now available algorithms that better handle anisotropic diffraction^31^. The resolution could thus be extended to 2.52 Å in the best direction, and 2.8 Å in the other two (Extended Data Table S1). The re-refined electron density maps improved substantially. Among other features, Q193_HK_ (Gln at position 193 of the HK) was now well defined in density, interacting with a water molecule that coordinates the Mg^2+^ cation (Fig. 1a). An additional water molecule is now clearly visible, hydrogen-bonded to the conserved T80_RR_ side chain (Fig. 1a), and well positioned to interact with the attacking water molecule for phosphoryl hydrolysis. Indeed, an unexpected electron density bulge coinciding with a Fourier difference peak was observed near the phospho-mimetic BeF_3_^−^ group, axially in line with the Be-OD54 bond to D54_RR_ (Extended Data Fig 2). This feature is consistent with a water molecule correctly placed to perform the nucleophilic attack, although limited data resolution precludes conclusive modeling.

**Figure 1:**
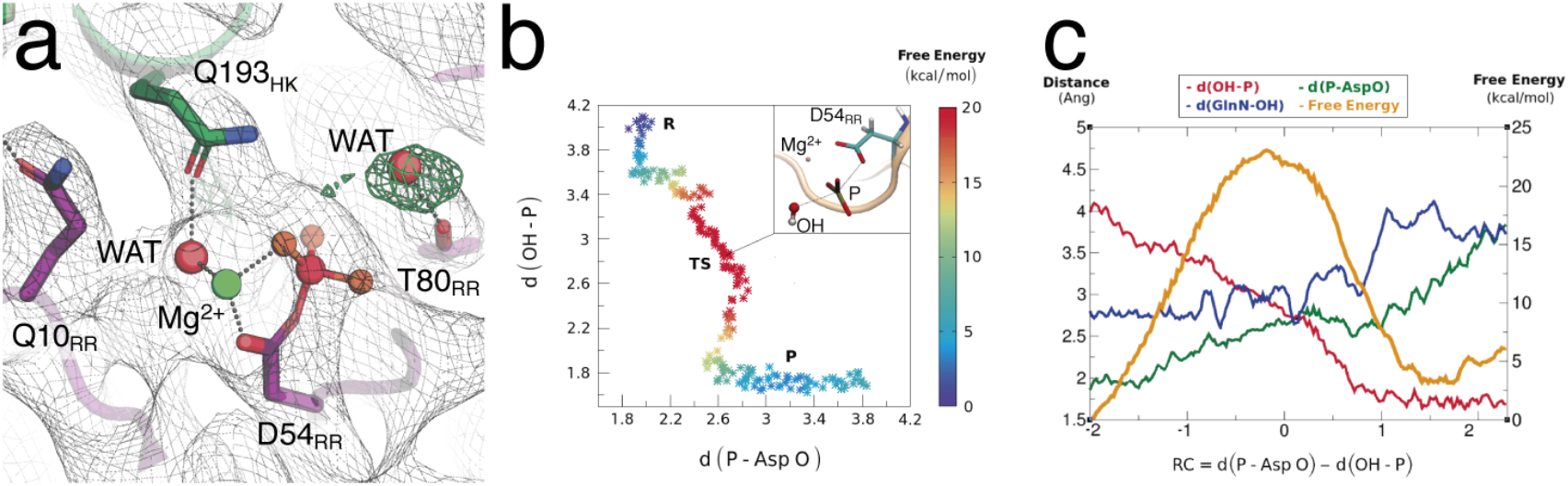
HK phosphatase-catalyzed reaction. a) Reaction center of the DesK-DesR complex in the phosphatase state. The gray mesh shows 2mFobs-DFcalc electron density map contoured at 1, and the green mesh shows positive peaks of the mFobs-DFcalc electron density contour at 3.5. Key residues are depicted in sticks. b) Bidimentional free energy diagram energy as a function of the bonds broken and formed for the phosphatase reaction. The distance between the phosphorus atom of the phosphoryl-moiety and the OE2 atom of D54_RR_ (P-Asp O) against the distance between the phosphorus and hydroxyl anion (P-OH) is shown in the plot as a way of describing the reaction path. Inset: QM region for QM-MM simulations of the phosphatase reaction. c) Free Energy profile (orange) and main interatomic distances as a function of the reaction coordinate for the phosphatase reaction, defined as the difference between the P-Asp O distance and the OH-P distance.

Our results showed that in order to simulate the P~Asp_RR_ hydrolysis reaction, the attacking nucleophile has to be a hydroxyl anion, and not a water molecule. Attempts to perform the reaction using a water molecule as the phosphate acceptor were unsuccessful, even if the transferring phosphate was concertedly probed as the proton acceptor. It must be stressed that the reaction center lacks a suitable residue that may act as a base to deprotonate the reactive water. Therefore, the hydroxyl must be formed in the bulk solvent. This observation is in accordance with the well-studied Ras/GAP proteins mediating GTP hydrolysis, where a solvent-assisted mechanism has indeed been put forward^32^.

The HK-mediated dephosphorylation reaction proceeds through a concerted nucleophilic substitution mechanism, as evidenced by the changes at the interactomic distance in the TS (transition state) zone (Fig. 1b and c), with an energy barrier of ~23 kcal/mol (Fig. 1b and c). The bond with the attacking water molecule occurs late in the TS (defined as the higher energy state of the reaction pathway), implying a dissociative mechanism with loose TS. The TS adopts the expected planar trigonal structure. Q193_HK_, which is bound to the Mg coordination sphere, remains hydrogen bonded to, and accompanying the acceptor OH^−^ along the reaction, only to release it after the TS is resolved, and the orthophosphate liberated as final product. The same QM/MM calculation was performed in the absence of Mg^2+^, displaying a much higher energy barrier. On the other hand, the calculated reaction without DesK, surprisingly showed a similar profile (Extended Data Fig 3), suggesting that additional mechanisms should be at play. Taken all the evidence together, the sidechain amide of the conserved Q193 on the HK’s α1 helix, appears to assist in the reaction by placing the hydroxyl anion in a catalytically ideal in-line attack position, which would otherwise be diverted by the action of the phosphoryl oxygens^33^.

### DesK:DesR phosphoryl-transfer follows a dissociative nucleophilic substitution mechanism

To test whether the difference in the distances between phosphoryl-acceptor and - donor residues correlates with a looser or tighter phosphoryl-transfer TS between phosphohistidine and aspartate, the free energy profiles of the reactions were computed using QM/MM-based steered molecular dynamics using either a HK:RR or a phosphorelay Hpt:RR type of complexes. First, to build the starting phosphotransferase-competent state of the DesK:DesR complex, the crystal structure of the DesKC_H188E_:DesR_REC_ ^1^ complex was used (PDB id:5IUK)^7^, into which the phosphorylated H188_HK_ was modeled by superimposing the crystal structure of phosphorylated wild type DesK alone (PDB id:5IUM)^7^. The reaction could be nicely simulated, exhibiting a low energy barrier (Fig. 2a, Table 1 and Extended Data Fig 4), and confirming that the starting point is a good representation of the phosphotransferase state.

**Figure 2:**
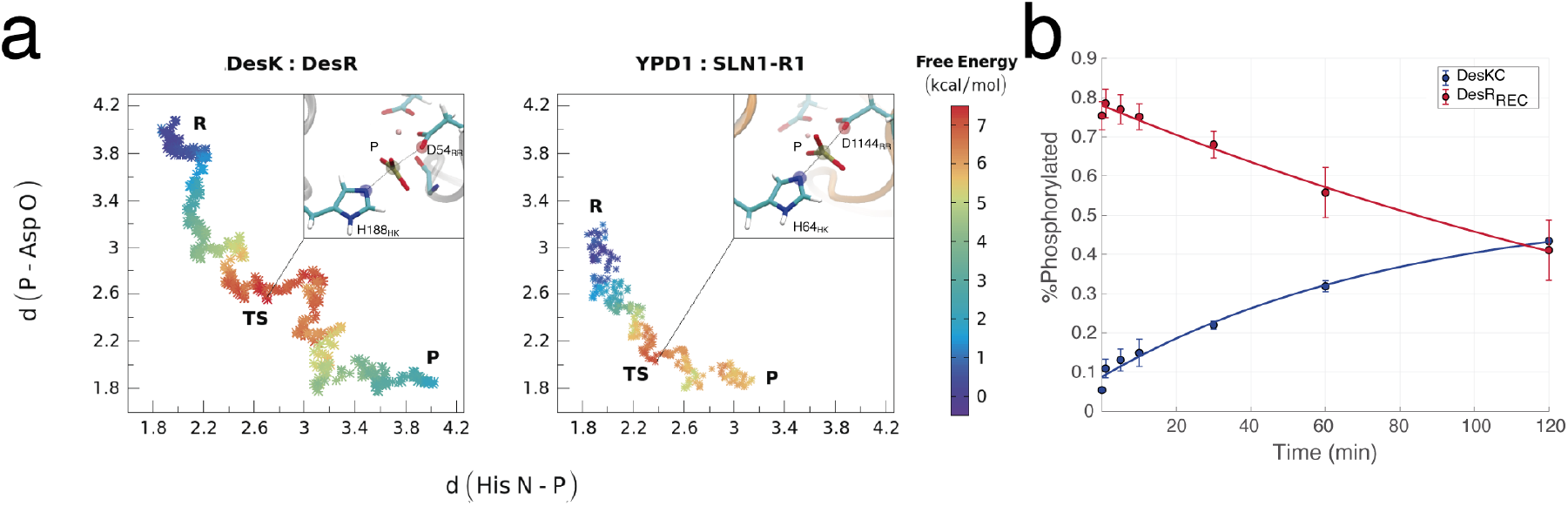
Phosphoryl-transfer reaction. **a)** Reaction diagram of the phosphoryl-transfer reaction. The left panel shows the bidimentional free energy diagram energy as function of the breaking and forming bonds of the DesK-DesR system (AspO-P vs HisεN-P). Right panel shows the Free Energy diagram for the reversible phosphorelay Sln1-Ypd1 systems. **b)** Phosphotransfer kinetics in the absence of Mg^2+^. Equimolar amounts of phosphorylated DesR_REC_ and DesKC were incubated and analyzed by densitometry from Coomasie stained Phostag SDS-PAGE. The solid lines shows the exponential fit of the phosphorylation degree of DesR-REC (red) and DesKC (blue).

**Table 1:**
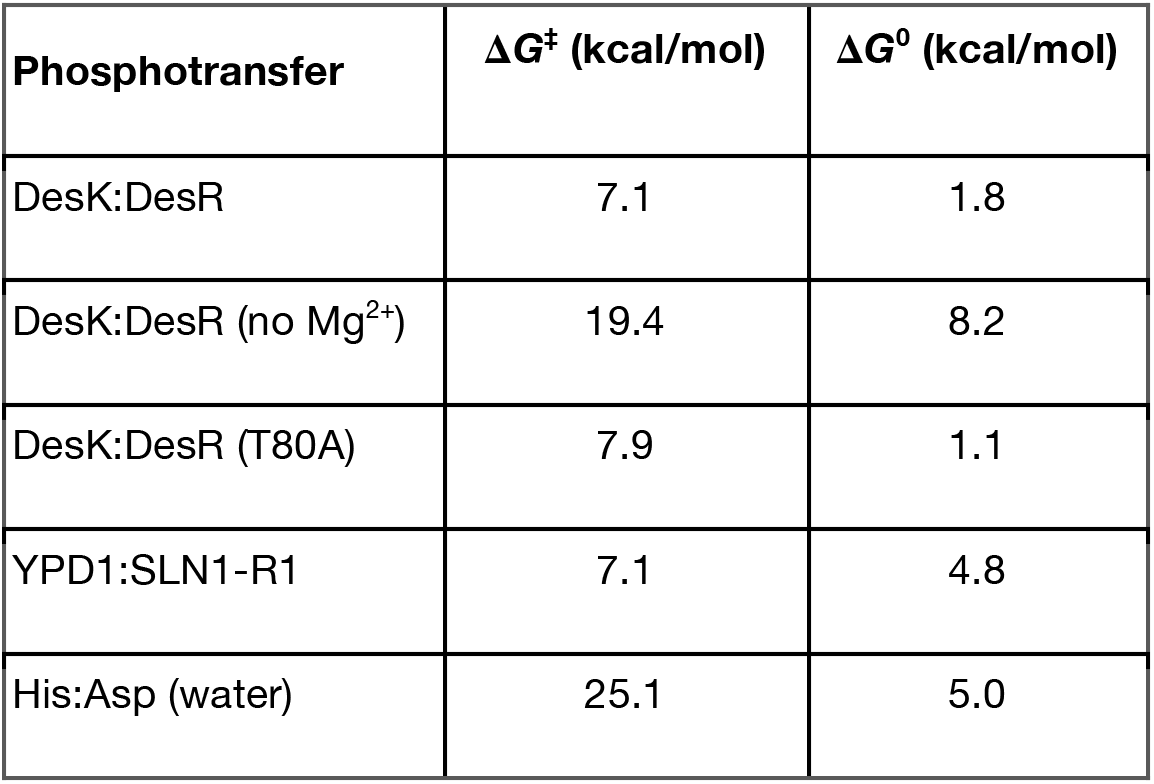
Free energy derived from QM/MM calculations

| Phosphotransfer | $\Delta G^\ddagger$ (kcal/mol) | $\Delta G^0$ (kcal/mol) |
| --- | --- | --- |
| DesK:DesR | 7.1 | 1.8 |
| DesK:DesR (no $Mg^{2+}$) | 19.4 | 8.2 |
| DesK:DesR (T80A) | 7.9 | 1.1 |
| YPD1:SLN1-R1 | 7.1 | 4.8 |
| His:Asp (water) | 25.1 | 5.0 |

The reaction starts with the phosphoryl moiety bound onto H188_HK_ and also establishing a hydrogen-bond with Thr80_RR_ (Extended Data Fig 4b). The latter hydrogen bond remained during the whole reaction. The TS shows the *en transfer* phosphoryl-group adopting a planar trigonal structure (metaphosphate). Remarkably, the TS zone shows a dissociative character. Although the reactive D54_RR_ got initially closer to the phosphoryl group (Fig. 2a), the O-P distance stopped decreasing at *ca*. 2.8 Å (*i.e*. the new covalent bond is not yet established) to then remains constant while the N-P bond is stretched. Only after the donor N-P bond increased to 3.1Å (*i.e*. bond cleavage) the D54_RR_ O-P bond was properly formed (Fig. 2a and Extended Data Fig 4b).

The role of the Mg^2+^ was assessed by calculating the free energy profile in the absence of the metal cation. This resulted in a much larger energy barrier and unfavorably positive reaction free energy (ΔG^*0*^), similar to the calculated phosphoryl-transfer reaction between isolated His and Asp residues in water (Table 1 and Extended Data Fig 4a). The positive ΔG^0^ for the isolated reaction is expected from a pure chemical viewpoint, as described in the introduction. Consistently, phosphoryl-transfer reactions using equimolar amounts of DesR_REC_~P and DesKC, in the absence of Mg^2+^ showed that the reaction proceeds slowly toward the His (Fig. 2b), spontaneously approaching maximum HK phosphorylation (*i.e*. hemiphosphorylation of the HK dimer)^29^. This indicates that the phosphoryl-transfer reaction itself is not completely abolished in the absence of Mg^2+^, but the equilibrium is highly displaced toward the His.

Secondly, to compare *bona fide* HK:RR complexes with those of phosphorelay pathways, similar free energy profile analyses were computed using the yeast osmosensor phosphorelay complex Ypd1:Sln1 (Fig. 2). In this case the reaction showed a more associative mechanism than for the DesK-DesR system, consistent with an initial shorter distance between the phosphoryl donor and acceptor residues, and a similarly small energy barrier of 7 kcal/mol (Table 1). The difference between the product free energy (phosphorylated Asp) and the starting point (phosphorylated His) is more positive in Ypd1:Sln1 as compared to the DesK-DesR system (Table 1 and Extended Data Fig 4a) and similar to the His:Asp reaction in water (Table 1). This is an important observation, consistent with our hypothesis of a correlation among associative phosphoryl-transfer reactions, shorter inter-atomic distances, and the chemical equilibrium displaced toward His phosphorylation, and *vice versa*. Altogether, our results thus provide strong evidence that the experimental crystal structure of the DesK_H188E_:DesR_REC_ complex is a good mimetic of the phosphotransferase state and that phosphoryl-transfer proceeds through a more dissociative mechanism or loose transition state in DesK:DesR than in the phosphorelay.

### Highly conserved D189_HK_ and T80_RR_ residues are not essential for phosphoryl-transfer

The configuration of the phosphotransferase active site places the highly conserved T80_RR_ at interaction distance with the covalently bound phosphoryl moiety of H188_HK_. This interaction was kept during the entire phosphoryl-transfer process in our QM/MM calculations. To further analyze the relevance of T80_RR_ in the reaction, the HK-mediated phosphorylation of T80A_RR_ (Fig. 3a) and T80S_RR_ (Extended Data Fig 5) mutants were measured. In the *in vitro* assays, phosphoryl-transfer and desphosphorylation of the RR are simultaneously taking place (biphasic curves). Intriguingly, DesK was able to phosphorylate these two DesR mutants, showing phosphoryl-transfer reaction comparable to wild type (Fig. 3b), thus indicating that T80_RR_ is not critical for the reaction. QM/MM calculations of the phosphoryl-transfer reaction using the T80A mutant, further suggest this residue is not essential (Table 1). Of note, T80A_RR_, but not T80S_RR_, was unable to be phosphorylated using acetyl-phosphate as a phosphoryl-donor (data not shown).

**Figure 3:**
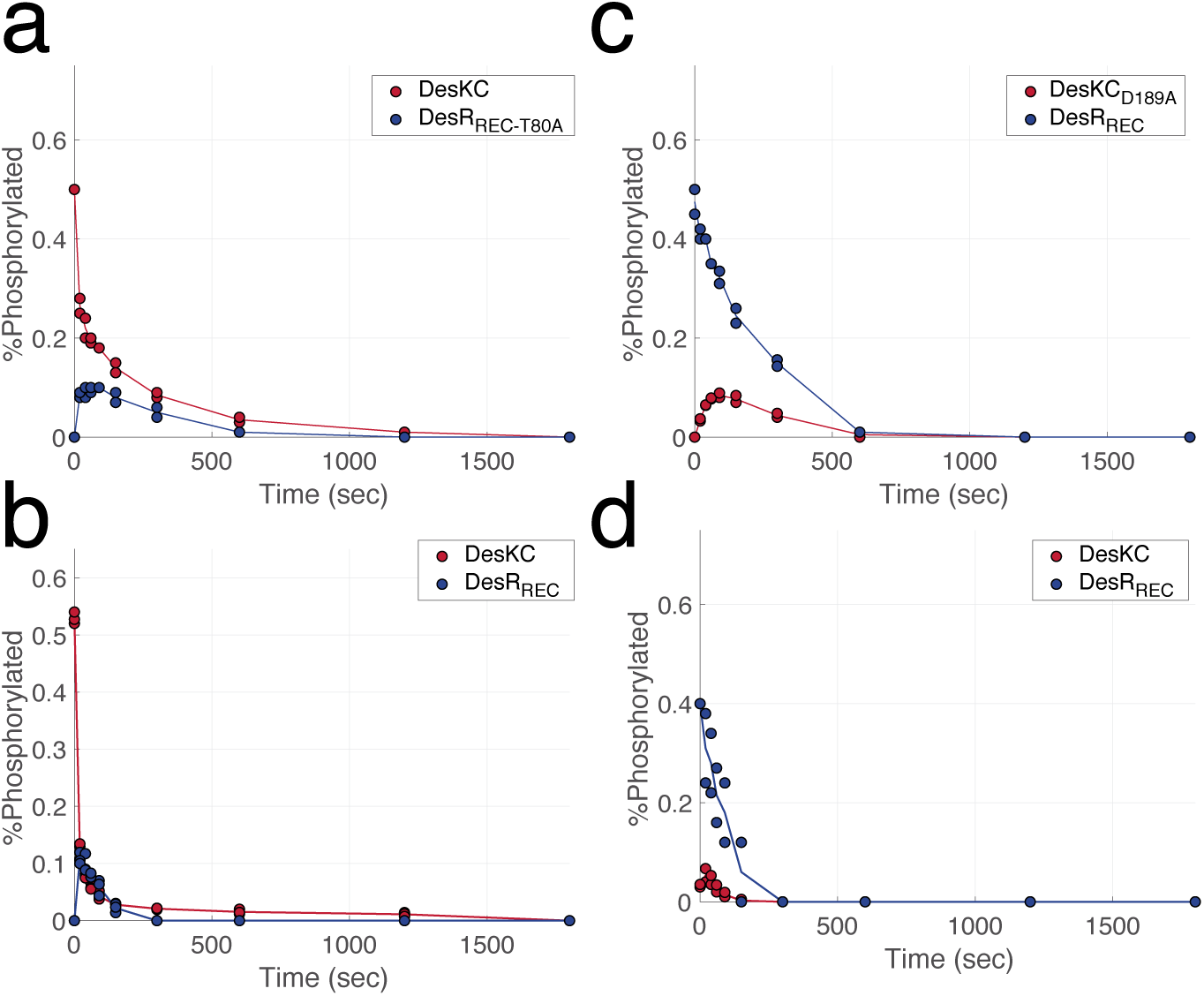
Analysis of highly conserved residues in the phosphoryl-transfer reaction. Assays were performed with DesKC~P in the presence of DesR_REC-T80A_ (a) or DesR_REC_ (b); or DesR_REC_~P in the presence of DesKC_D189A_ (c) or DesKC (d). Degree of phosphorylated of DesKC (red) and DesR_REC_ mutants (blue) was analyzed *in vitro* by densitometry of Coomasie stained PhosTag SDS gels. Two independent experiments are shown for each reaction.

Concerning D189_HK_, it can be predicted that this highly conserved acidic residue at position H+1 (one residue C-terminal to the phosphorylatable His), is not involved in the phosphoryl-transfer reaction. This is in contrast with the critical role that this residue plays in subtracting a proton from the His δN, increasing the nucleophilicity of the His in the autophosphorylation reaction^10,13,34,35^. In the phosphotransferase state, due to the rotameric configuration of H188_HK_, the δN is oriented in a way that cannot interact with D189_HK_. Consistent with this view, reverse phosphoryl-transfer of a D189A_HK_ mutant was not abolished (Fig. 3c).

### Phosphoryl-transfer reversibility is modulated by amino acid substitutions on the RR

QM/MM calculations suggested that, although a looser TS could be associated with a shift in the equilibrium towards phosphorylation of the RR, when compared to the more associative mechanism of the phosphorelay system, the ΔG^*0*^ of the reaction was still positive. Thus, on thermodynamic grounds, the transfer reaction appears to always favor the phosphorylation of the HK. In this scenario, the following question emerges: Have TCSs evolved additional mechanisms to deal with this energetic uphill? Three different mechanism could be at play: i) dimerization of the phosphorylated form of DesR could shift the equilibrium towards its phosphorylated state, especially considering that its α1α5 surface is used for both RR dimerization and DesK-interaction; ii) phosphorylated DesR might exhibit decreased affinity for DesK when compared to the unphosphorylated species; and/or iii) in the absence of the sensor domain, a shifted conformational equilibrium of DesK towards the phosphatase-competent state would reduce the amount of available phosphorylatable His, given that it is occluded inside the **D**imerization and **H**istidine **p**hosphotransfer domain (DHp)^7^. Since the interaction surfaces between DesK and DesR, comparing the HK’s phosphatase- and phosphotransferase-competent states, are very similar^7^, as are the conformations of bound DesR, the second option of affinity changes seems unlikely. Also, if DesR dimerization were to sequester the phosphorylated monomeric DesR species out of equilibrium, this should also preclude DesK-mediated dephosphorylation, which is clearly not the case. Thus, although dimerization could contribute to the observed P~His_HK_→Asp_RR_ directionality, it seems not be the main driving force.

We reasoned that if the phosphatase/kinase equilibrium of DesKC is shifted towards the former state, mutating residues that selectively affect DesR’s interaction with the phosphatase-competent state of the kinase only would induce a shift in phosphoryl-transfer reversibility. To test this hypothesis two DesR mutants, R84A_RR_ and Q10A_RR_, were tested. Residue R84_RR_ interacts specifically with DesK’s D189_HK_^7^, and due to the gear-box mechanism^9^, the R84_RR_:D189_HK_ interaction is disrupted in the phosphotransferase state through the rotation of the DHp domain α-helices (Fig. 4a). The R84A_RR_ substitution significantly reduced the dephosphorylation of the P~DesR compared to wild type (Fig 3d), but the phosphoryl group still accumulates in DesR (Figure 4b).

**Figure 4:**
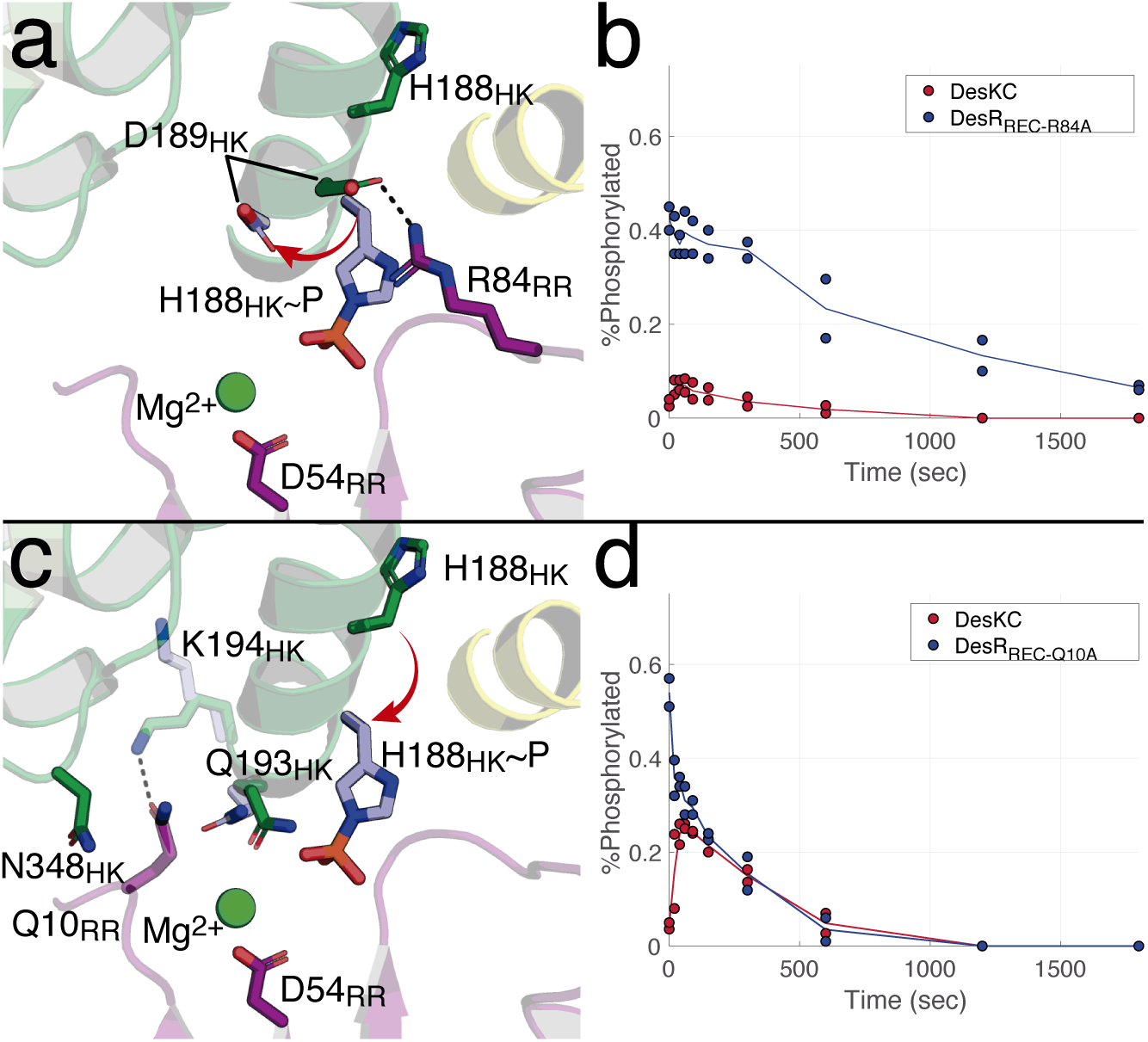
Modulation of phosphoryl-transfer reversibility. (a) Structural comparison between the phosphotransferase (residues depicted in light blue) and phosphatase (in green) complexes highlighting the interaction established by R84_RR_. (b) Phosphoryl-transfer assay showing the distribution of the phosphoryl-moiety between DesKC and phosphorylated DesR_REC-R84A_. (c) Similar view as in (a), showing Q10_RR_ inserted in a pocket created at the ATP binding domain:DHp interface, and interacting with K194_HK_. (d) Phosphoryl-transfer assay of DesKC and phosphorylated DesR_REC-Q10A_.

On the other hand, residue Q10_RR_ is inserted in a pocket generated at the interface between the DHp and ATP binding domain (Figure 4c), a hallmark of the phosphatase state^36^. Remarkably, a Q10A_RR_ mutant showed a significant increase in phosphoryl-transfer reversibility, with the phosphoryl moiety more evenly distributed between the kinase and the regulator (Fig. 4d). Q10_RR_ is far from the phosphorylation site and was previously highlighted as highly covariant with Q193 on the HK^7^. To rule out that transfer reversibility of the Q10A_RR_ mutant is caused by a different positioning of DesR in the phosphotransferase complex, potentially shortening the His_HK_-Asp_RR_ distance, the crystal structure of this complex was solved at 3.4 Å resolution. The structure clearly indicated that the REC domain of DesR remains in the same position compared to the wt (Extended Data Fig 6 and Table S1), especially not altering the His-Asp distance. The observed change in phosphoryl-transfer reversibility is thus based on other reasons, and directionality can be shifted by mutations at the response regulator that are far from the phosphorylation site.

### A shifted HK conformational equilibrium minimizes phosphoryl-transfer reversal

To better understand the key determinants of information transmission, a systems biology and integrative approach was followed. Taking advantage that the different functional states of DesK can be trapped^37^, an experimental dataset comprising 770 measurements of the level of phosphorylation level in each protein (HK or RR) was generated at different time points along the phosphoryl-transfer reaction assays. Two types of reactions were analyzed, either starting with the phosphorylation of the HK (Fig. 5c,d,e,f and g) or the RR (Fig. 5h and Extended Data Fig 7a,b,c,d, e and g). To this end we used DesKCwt, which lacks the transmembrane sensor domain, and was previously shown to be deregulated^9,38^. DesKC_Q193A_ was used to evaluate phosphoryl-transfer with no confounding phosphatase activity. Q193_HK_ (H+5 position) is well conserved in all HisKA_3 HKs and critical to keep phosphatase activity^39,40^. DesKC_DEST_ was also included, since this is a mutant that shifts the conformational equilibrium of DesK towards the auto-kinase active state, by destabilizing the N-terminal coiled-coil^8^. DesKC_H188E_ and DesKC_H188V_ are constructs that trap DesK in the phosphotransferase^7,9^ and phosphatase state^38^, respectively. In addition, wild type DesR_REC_ and the point mutant DesR_REC-Q10A_, were also tested. The analyzed reactions confirmed several experimentally observed behaviors, such as the decrease in phosphatase activity of DesKC_Q193A_ (Fig. 5e and Extended Data Fig 7c), or the increased phosphoryl-transfer reversibility of DesR_REC-Q10A_ (Fig. 5d and f and Extended Data Fig 7b and d).

**Figure 5:**
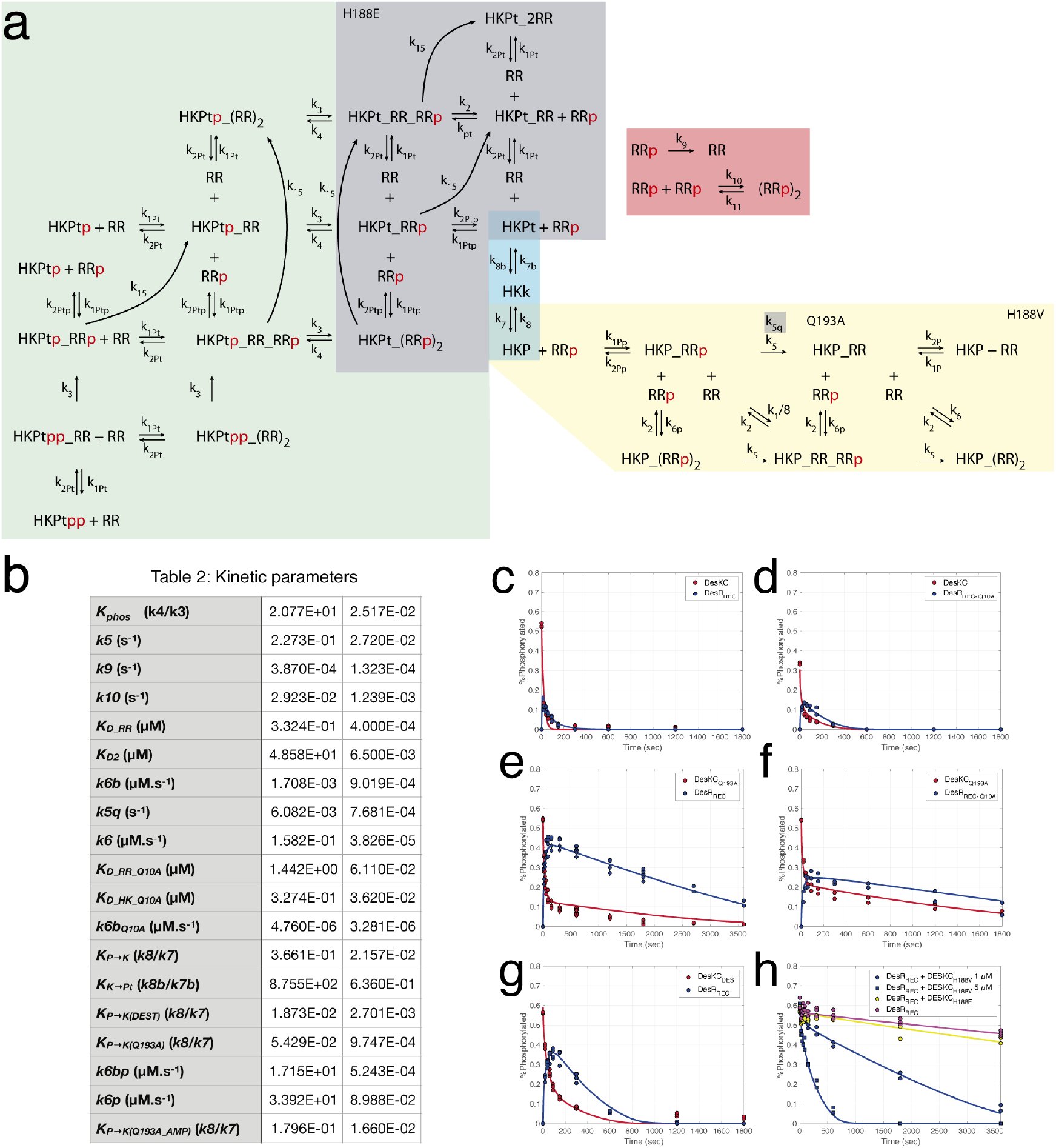
Phosphoryl-transfer assay dataset and model fitting. (a) Schematic representation of the kinetic model. Yellow shaded box are all the reactions involved in the HK-catalyzed dephosphorylation reaction (H188V module). The red box includes the dimerization and autodephosphorylation of the RR module. The gray box corresponds to the H188E module, which includes RR binding to the phosphotransferase, without phosphoryl-transfer reaction. Green box describes the phosphoryl-transfer reactions, and in cyan the HK conformational transitions are represented. (b) Parameters of the best fitted model. (c to g) Phosphoryl-transfer assays. The reactions were performed using either DesR_REC_ (c, e and g) or DesR_REC-Q10A_ (d and f) and started with phosphorylated wild type DesKC (c and d), DesKC_Q193A_ (e and f) or DesKC_DEST_ (g). (h) Phosphatase assay incubating phosphorylated DesR_REC_ alone, with DesKC_H188V_ or DesKC_H188E_. The continuous trace (red and blue lines) depicts the prediction of the best-fitted model.

Different kinetic models were constructed and optimized, taking into consideration the ensemble of available structural and biochemical data. The best-fitted model includes 19 free parameters (depicted in Fig 5a) and converged to a unique solution (Fig 5b). Residuals to test the significance of the differences among alternative models were rigorously monitored (see methods).

Several salient conclusions can be drawn from the optimized integrative model: i) the equilibrium constant of the phosphotransferase reaction is directed towards His_HK_ phosphorylation, as expected from our QM/MM calculations (*K_phos_* (k4/k3) = 20.7, Table 1); ii) the kinetic rate constant (*k5* = 0.227 s^−1^) of the HK-mediated phosphatase activity is 3 orders of magnitude higher than the intrinsic dephosphorylation rate of the RR (*k9* = 3.87E-4 s^−1^, autodephosphorylation); iii) the Q193A_HK_ substitution reduces the phosphatase activity by 2 orders of magnitude (*k5q* = 6.082E-3 s^−1^), but the activity is not completely abolished; iv) the dissociation equilibrium constant of P~RR dimerization is in the low μM range, and the Q10A_RR_ amino acid replacement increases this K_D_ by a factor of 4 (*K_D RR_Q10A_* = 1.442 μM); v) the phosphatase/auto-kinase conformational equilibrium of the HK is only slightly displaced towards the auto-kinase state; vi) Q193A_HK_ and DesKC_DEST_ variant shift the phosphatase/auto-kinase equilibrium toward the auto-kinase state; and, interestingly, vii) the auto-kinase/phosphotransferase equilibrium is highly shifted towards the auto-kinase state, preventing phosphoryl back-transfer.

The kinetic model also predicts that DesKC will interact with DesR mostly in a phosphatase conformation. Isothermal titration calorimetry (ITC) data indeed confirmed this hypothesis. DesKC associated to DesR with a 2:2 stoichiometry (two RR molecules per HK dimer) as had been previously observed using a phosphatase-trapped mutant^7^. Moreover, the relatively similar K_D_ values of the two independent binding sites was consistent with a symmetric architecture as evidenced in the phosphatase configuration (Extended Data Fig 8). The phosphotransferase configuration, due to its asymmetry, exhibits substantially larger differences in the K_D_ values^7^. The kinetic model allows us to make further predictions, such as a shift of the dimerization equilibrium in the DesR Q10A_RR_ mutant, driving it towards the monomeric species (higher dissociation constant of dimerization K_DRRQ10_ compared to wild type). Size-exclusion chromatography of phosphorylated DesR-REC_Q10A_ indeed showed disturbed dimerization for this mutant compared to the wild type protein (Extended Data Fig 9). Taking all the evidence from the model together, we conclude that phosphoryl-transfer directionality is dictated by: i) the auto-kinase/phosphotransferase conformational equilibrium, ii) the ratio between the forward and reverse phosphoryl-transfer kinetic rates, and iii) the dimerization of the P~RR.

## Discussion

### A subtle displacement of the RR in the HK:RR interaction minimizes futile cycles

Signal transduction pathways that use phosphorylation as a means to transmit information use cellular energy by consuming ATP. In this context a key question is: Have molecular machines evolved to minimize unwanted energy waste? There are examples where energy dissipation is biologically relevant, for instance to deal with noisy signaling and allow for adaptation^41^. In the case of TCSs, HK bifunctionality confers robustness to the signaling process^42,43^, but raises the question of how tightly it must be regulated to avoid futile cycles and energy waste. In the present work we described from first principles the HK-mediated dephosphorylation reaction of P~RR. The crystal structure of the DesK:DesR phosphatase complex shows well-defined density of key residues. That the phospho-mimetic BeF_3_^−^ group bound to D54_RR_ shows an unexpected bulge in the electron density suggest that instead a MgF_3_^−^ (magnesium fluoride) could be present as mimetic of the transition state ^44^ (Extended Data Fig 2). In any case, the architecture of the reaction center in the X-ray structure is perfectly consistent with the QM/MM calculations. The HK favors hydrolysis of the P~Asp by organizing the active site within the RR, assisting with its glutamine in position H+5 in the correct placement of the attacking hydroxyl anion. The interaction between the HK in the phosphatase conformation and the phosphorylated RR promotes the opening of the RR active site, exposing the phosphorylation site^7,28^. According to our kinetic model, the dephosphorylation of the RR is accelerated by the HK by 3 orders of magnitude compared to the intrinsic autodephosphorylation reaction. Moreover, in the absence of the conserved Q193_HK_ (Q193A substitution), the reaction is 2 orders of magnitude slower compared to wild type HK. This modest acceleration and the effect of Q193 mutation is consistent with the proposed role of Q193_HK_. The QM/MM calculations of the dephosphorylation reaction in the presence or absence of the HK are similar, implying that the acceleration promoted by the HK is likely gained through lowering the entropy of the Michaelis complex. Analogous mechanisms have been put forward after extensive studies of the Ras/Ras-like small GTPase family of proteins. In the case of Ras, a GTPase activating protein (GAP) accelerates the reaction by several orders of magnitude^45^. The active site is very similar to the reaction center conformed between the HK and the RR in TCSs (Extended Data Fig 10). GAP binding to Ras induces a significant loss of entropy of the Gln, which was proposed to be critical in the GTP hydrolysis reaction mechanism^32^.

When the HK is stabilized in its phosphotransferase conformation, to avoid futile cycles the dephosphorylation of P~RR must be minimized. From a structural point of view, even though differences in the HK active site are observed comparing its phosphatase and phosphotransferase conformations, the position of Q193_HK_ is unchanged and the conformation of the RR is almost identical^7^. Then, why is there no significant phosphatase activity? An explanation can be uncovered by superimposing the DesK:DesR phosphatase and phosphotransferase complexes. There is a small but significant shift (approximately 1 Å) in the position of the RR. This movement, in the phosphatase state, brings the phosphoryl-group and the Mg^2+^, with its entire coordination sphere, closer to Q193_HK_ (Extended Data Fig. 11). Hence, Q193_HK_ comes now within interaction distance with a water molecule of the Mg^2+^ coordination sphere, properly placed to position the attacking hydroxyl anion. The ~1Å shift of the RR is driven by a set of interactions between the REC domain and the HK ATP binding domain, that are only available in the phosphatase state. In contrast, in the phosphotransferase state, Q193_HK_ is not ordered, as evidenced by the electron density, which is not well defined (Extended Data Fig 11). Collectively, our data suggest that interactions involving secondary interfaces in the phosphatase state, more precisely between the REC and ATP binding domain, guide the correct placement of the RR to favor dephosphorylation. In the absence of this interaction, the RR is held farther apart and thus not promoting the phosphatase reaction.

### Insulation of HK activities as an information-driving mechanism

The higher energy of the mixed anhydride O-P bond on the P~Asp compared to the N-P phosphoramidate in P~His^25^, which is in turn higher than the anhydride O-P bond of ATP^46^, imposes an uphill energy barrier that the system must overcome to promote RR phosphorylation. A shift in the phosphoryl-transfer reaction equilibrium according to the distance between nucleophile and leaving group amino acids could be possible, considering the position of the Mg^2+^ cation. The relative position of the cation changes with respect to the HK, in different complex geometries. Our QM/MM calculations of phosphoryl-transfers along looser or tighter TS, suggest that changing the distance between the donor and acceptor residues can indeed setup different ΔG of the reaction. Nevertheless, the ΔG is always positive in the P~HK→RR direction, so that additional mechanisms are at play.

The structural model of the DesK-DesR system (Fig. 6a) suggests that the proper accumulation of P~RR could be reached by: i) P~RR dimerization; ii) a reduction in binding affinity to the HK when the RR is phosphorylated; and iii) through regulation of the conformational equilibrium of the HK, making the phosphotransferase conformation unlikely. According to the integrative kinetic model, the binding affinity of the HK to the RR in phosphorylated and non-phosphorylated states of the latter, and either in the phosphatase or phosphotransferase state of the HK, is unchanged. This is consistent with all the experimental information, in particular the similarity of the phosphatase and phosphotransferse binding areas at the HK:RR interface. Moreover, a decrease in the affinity to the phosphorylated RR would also imply an inconsistent scenario where the HK in the phosphatase state would be unable to interact with it. P~RR dimerization does compete with HK binding, and the observed effect of the Q10A_RR_ mutant shows that disturbing this equilibrium indeed results in a more reversible phosphoryl-transfer. However, too tight a dimerization would also preclude HK-dependent and independent dephosphorylation, which are constraints that need to be considered.

**Figure 6:**
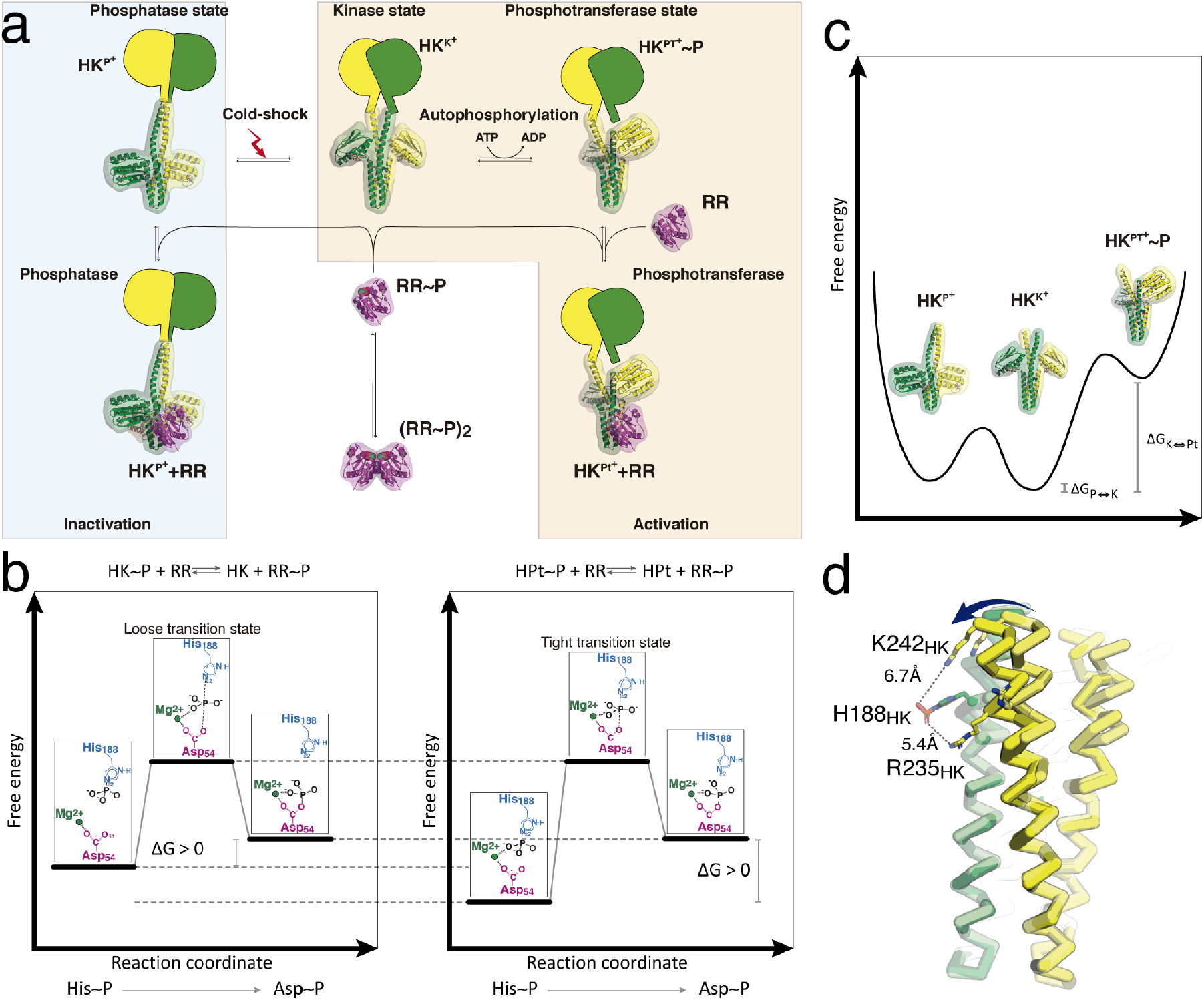
DesK-DesR structural and kinetic model (a). b) Diagrammatic representation of the free energy transition of the phosphoryl-transfer reaction of a more loose (left panel) or tight (right panel) TS and the role of the Mg^2+^. A shorter His-Asp distance places the phosphoryl moiety at interaction distance to the coordinated Mg^+2^ at the beginning of the reaction, thereby stabilizing the initial state c) Schematic representation of the conformational equilibrium of DesKC in the absence of phosphorylation, indicating the higher energy of the phosphotransferase state. d) Proposed role of R235_HK_ and K242_HK_ in sensing the phosphorylation state of the His.

From our kinetic model, the parameters that contribute the most to the directionality are: the ratio between the kinetic rates of the phosphotransferase along its forward and reverse directions, and the conformational equilibrium between the HK’s auto-kinase and phosphotransferase states (Extended Data Fig 12). By moving the reactive His closer or farther apart in the reaction center, the ΔG might be modulated due to the key contribution of the Mg^2+^ cation at the beginning of the reaction, stabilizing the phospho-His in case the distance is short enough (Fig. 6b). This is in agreement with the ΔG calculated for the isolated His to Asp phosphoryl-transfer reaction in water (Table 1), suggesting that in an associative mechanism the influence of the Mg^2+^ in shifting the equilibrium towards Asp phosphorylation is lower. The solvent-occluded position of the His inside the DHp core of the HK in the phosphatase conformation^7^ should minimize reversibility as well. Mutants that shift the phosphatase/auto-kinase equilibrium should be more reversible. We did not observe such effect when we tested DesK_DEST_ (Fig. 5 and Extended Data Fig 7). On the other hand, our finding that the equilibrium between the auto-kinase and phosphotransferase conformations was shifted against the latter, which is the only competent state able to participate in the phosphoryl-transfer reaction, also explains why the His is not accessible anymore (Fig. 6c). From a structural point of view it appears that phosphorylation favors the phosphotransferase state by triggering a conformational rearrangement. The highly conserved R235 and K242 in HK helix a2 are well positioned to interact with the phosphoryl-group on the reactive H188. This interaction likely pulls the a2-helix toward the a1-helix, breaking the symmetric organization of the DHp and favoring the phosphotransferase state (Fig. 6d). A tight non-covalent interaction between the guanidinium group of the Arg with the phosphate^47^ could provide the necessary energy to promote this somehow unfavorable conformational transition.

The unexpected conformational equilibrium uncovered by our kinetic model also indicates that the system is tuned to maximize signal transmission efficiency. In other words, in the presence of the specific signal, HK activation promotes a shift in the phosphatase/kinase conformational equilibrium towards the latter. However, depending on the kinase/phosphotransferase conformational equilibrium and intracellular protein concentrations, HK autophosphorylation might be inhibited by its the interaction with the RR in the phosphotransferase state. Our kinetic model predicts that there is a narrow window of the kinase/phosphotransferase equilibrium in which the system can accumulate phosphorylated RR, and requires that in the absence of phosphorylation, the phosphotransferase state of the HK is inaccessible (Extended Data Fig 13).

Taking all together, we conclude that the sensor domain controls the on/off switch by changing the auto-kinase/phosphatase transition. But, the auto-kinase/phosphotransferase equilibrium guarantees that the information cannot go backwards. Finally, our findings might also be extrapolated to phosphorelay pathways, in which different mechanisms could allow these systems to be more or less reversible, by tuning the Asp-His distance, the dimerization of the phosphorylated effector RR, or additional elements like local physicochemical properties in the immediate surroundings of the P~Asp that were not considered in this work.

## Methods

### Classical (MM) Simulations

All molecular dynamics simulations were performed using the AMBER suite package of software^48^. Classical force field parameters for aminoacids were obtained from the ff14SB force field, whereas force field parameters for phosporylated residues (phospo-Asp and phospho-his) were generated with the Antechamber module of the AMBER suite, after geometry optimization and RESP charge derivation at the HF/6-31G* level with Gaussian g09, revision D.01^49^. For each simulation, the equilibration protocol consisted of 200-cycle runs of minimization with a 100 (kcal/mol)/Å^2^ restraint constant applied to the protein in order to relax the solvent structure, followed by a 1000-cycle energy optimization in which the restraint was removed to avoid initial unfavorable contacts. The system was then slowly heated to 300 K during a 1 ns simulation, with the Berendsen thermostat. Finally, pressure was equilibrated at 1 atm over 1 ns, to let the system reach the proper density. For all simulations we used the periodic boundary condition approximation, with the Ewald summation method with a 10 Å cutoff for nonbonded interactions, and the SHAKE algorithm for all hydrogen-containing bonds. Final production 10 ns MD simulations were performed at 300 K using the Langevin thermostat and a 2 fs time step, from which the last structure was selected for QM-MM simulations.

### Hybrid (QM-MM) Simulations

All DFT QM/MM calculations in this work were performed with the SANDER(AMBER) program and the QM(DFT)/MM implementation called LIO^50^ using the PBE functional and a DZVP gaussian basis set. All relevant parts of the computation of LIO are ported to GPU, obtaining an improvement on the performance, including exchange and correlation, Coulomb, and QM/MM coupling terms^51,52^.

For the phosphotransfer reactions, the QM region consisted of 70 atoms (Extended Data Fig 14), including both donor and acceptor amino acids (His/Asp), the magnesium ion and its whole coordination sphere (sidechain of D9_RR_ and D54_RR_, backbone carbonyl of E56_RR_, two water molecules, and the phosphate) and a total of 8 H-link atoms at each corresponding boundary. For the phosphatase reaction the QM region is the same but with the donor His replaced by a hydroxyl anion.

### Multiple Steered Molecular Dynamics protocol

To study both the phosphotransfer and the phosphatase reaction mechanisms, we used a multiple steered molecular dynamics (MSMD) strategy, combined with Jarzynski’s relationship to determine the corresponding Free Energy Profiles. This strategy has already been shown to be useful in previous works from our group for phosphoryl transfer reactions^35,53,54^. Briefly, in MSMD the system is driven “multiple” times along the selected reaction coordinate under nonequilibrium conditions, by applying an external force, and for each individual trajectory, the work performed by the external force is computed. Finally, multiple works are exponentially averaged using Jarzynski’s relationship to obtain the free energy profile. In the present work we first performed 5 ns of conformational sampling at the reactive/product equilibrium geometries using standard QM/MM Molecular Dynamics. This was followed by 10 independent MSMD simulations (starting from corresponding initial structures each separated by 500 ps) which were run for approximately 2 ps using a 0.001 ps time step, resulting in a pulling speed of 2 Å/ps. The force constant used was 200 kcal.mol^−1^.Å^−2^. All the reaction mechanisms were simulated in both forward and reverse directions, and in each case the reported free energy profile corresponds to the optimal combination of both, as usually done when applying this strategy. The reaction coordinate for each profile was always the difference between the donor atom to P and P to acceptor (attacking atom) distances (See Extended Data Table S2 for values)

### Cloning, Protein Expression and Purification

Cloning and protein purification were performed as described previously^7,9,28^. Briefly, plasmid pQE80_DesKC_DEST_ was generated by subcloning DesKC_DEST_ from pHPKS/Pxyl-desKDEST^8^ into pQE80-DesR through PCR amplification using primers DesK_BamHI_F (CACGGATCCAGCAAGGAGCGCGAACGACTTG) and DesK_SalI_R (TCCTGGTCGACTTA TTTTGAATTATTAGGAATTGC), BamHI and SalI digestion, and ligation. pQE32-DesKC_Q193A_ was constructed using primers DesKQ193A (GATACGCTTGGGGCAAAGCTTT CTC) and DesK_Rev (GAATTATTAGGAATTGCCATGGTAAGCTTGGTC) by RF cloning^55^. pQE80_DesR-REC_Q10A_ was generated by similar procedure using DesR_Q10A_F (5’-GTATATTTATTGCAGAAGATGCGCAAATGCTGCTGG-3’) DesR-REC_R (5’-GCTTCGCTGTATAAGTCCTCCATCAG-3’). Recombinant proteins DesKC, DesR-REC and derived mutants were expressed as N-terminally His6-tagged fusions in *E. coli* strain TOP10F’. The last purification step was a size exclusion chromatography (HiLoad 16/60 Superdex 75 preparation grade column; GE) equilibrated with 20 mM Tris-HCl pH 8.0, 0.3 M NaCl (SEC buffer). All proteins were concentrated to ~10 mg/mL and stored at −80°C.

### Autodephosphorylation, Phosphotransferase and Phosphatase Assays

Phosphorylation of DesKC was performed by incubating purified DesKC with 10 mM ATP and 10 mM MgCl_2_ for an hour at 24°C in SEC buffer. The reaction mix was then loaded in a Superdex75 10/300 column (GE Healthcare) equilibrated in with a solution containing the SEC buffer. To obtain phosphorylated DesR_REC_ and DesR_REC-Q10A_, each pure protein (~600 μM) was autophosphorylated using 50 mM acetyl phosphate and 30 mM MgCl_2_ for an hour at 24°C, in the same buffer. Reactions were stopped by adding EDTA (50 mM), and loaded into a Superdex75 10/300 column (GE Healthcare) equilibrated in buffer SEC. The peak corresponding to the dimeric species was selected for further analysis.

The phosphotransfer assay was started by incubating P~DesKC (wild type or mutant) at a concentration of 26 μM (concentration of the monomeric species) with equimolar amounts of DesR_REC_ (wild type or mutant) in reaction buffer (20 mM Tris-HCl pH 8.0, 0.3 M NaCl and 5 mM MgCl_2_) for 30’ at 24°C. At different time points the reactions were stopped by adding SDS-PAGE sample buffer with 2.5 mM of DTT. Excess DTT was blocked with 40 mM iodo-acetamide and loaded in a Phos-tag SDS-PAGE, as described before^28^. Coomassie blue-stained gels were scanned and quantification was done by densitometry using ImageJ^56^.

Auto-dephosphorylation of P~DesR_REC_ (wild type or mutants) was performed as above, incubating P~DesR_REC_, at different protein concentrations, in Reaction buffer at 24°C. On the other hand, DesK promoted DesR dephosphorylation was done by incubating 26 μM P~DesR_REC_ in the presence of DesKC (wild type or mutants) at 26 μM (except for DesKC_H188V_ that different concentrations were tested) in Reaction buffer at 24°C. Reactions were stopped at different time points and loaded in a Phos-tag SDS-PAGE as described above.

### Protein Crystallization, Data Collection and Model Building

The DesKC_H188E_:DesR_REC-Q10A_ complex was prepared by mixing 300 μM DesKC_H188E_ and 165 μM DesR_REC-Q10A_ in a buffer containing Tris-HCl 20 mM pH 8.0, 0.3M NaCl, 20 mM MgCl_2_ and 5 mM AMP-PCP (non-hydrolysable analogue of ATP). The complex crystallized in a mother liquor containing 20% (w/v) PEG 3350 and 0.35 M tri-potassium citrate^7^. Protein drops were setup by mixing 2 μL of protein plus 2 μL of mother liquor. Cryo-protection was achieved by quick soaking in 20% (w/v) PEG 3350, 0.35 M tri-potassium citrate, 5 mM AMP-PCP, 25% (v/v) glycerol, and 20 to 150 mM MgCl_2_ + 5 mM BeF_3_^−^.

Single crystal X-ray diffraction was performed in a copper rotating anode home source (Protein Crystallography Facility, Institut Pasteur de Montevideo). Diffraction data was processed with autoProc^57^ and structure was solved by molecular replacement using each molecule of 5IUK^7^ as search probe in Phaser^58^. We reprocessed the X-ray diffraction data from 5IUN by merging two dataset from the same crystal in autoPROC. Model building was done in Coot^59^ and refinement in Buster^60^. Validation was done throughout and towards the end of refinement using MolProbity tools^61^. Visualization of protein models and structural analyses and figure rendering were performed with Pymol^62^. Software for data processing, structure determination and analysis was provided by the SBGrid Consortium^30^.

### Analytical Size Exclusion Chromatography

To analyze phosphorylation-triggered dimerization of Q10A_RR_ substitution, recombinant purified proteins dimeric phosphorylated DesR_REC_ and DesR_REC-Q10A_ Dimeric species were obtained as described above and degree of phosphorylation for both proteins was determined by Phostag SDS-PAGE. Then 100μL of 100 μM protein, with a degree of phosphorylation of 60%, was loaded in a Superdex75 10/300 column (GE Healthcare) equilibrated with a buffer containing 20 mM Tris-HCl pH 8.0, 0.3 M NaCl and 10 mM MgCl_2_ and run at 0.5 mL/min.

### Kinetic Model of the DesK-DesR System

Several kinetic models were constructed taking into consideration biochemical, structural and biophysical information from previous work^7,9,28^. We considered models with two or three functional states of DesK, in the latter case assuming the kinase state to be different from the phosphotransferase state. The autophosphorylation reaction was not included in the model, since all the reactions were performed in the absence of ATP or ADP. In addition, we tested models in which the binding of DesK might occur with identical or different affinity when DesR is phosphorylated or not. We also tested a simplified model where we discarded the second binding site for DesR in the phosphotransferase state of DesK. The best-fitted model is depicted in Fig. 5. Each model was constructed as custom scripts in Matlab R2020a (MathWorks Inc.), consisting in a set of differential equations that describes each reaction, conformational rearrangement (like kinase/phosphotransferase transition) or interaction between proteins. The model equations were solved numerically using the ode15s solver. Parameters were globally optimized by minimizing the sum of squared residuals (SSR) of the full dataset with the simplex search method as implemented in fminsearch with boundaries (John D’Errico (2021). fminsearchbnd, fminsearchcon (https://www.mathworks.com/matlabcentral/fileexchange/8277-fminsearchbnd-fminsearchcon), MATLAB Central File Exchange). Given the complexity of the parameter space, an exhaustive search was performed, starting from 800000 different initial conditions, uniformly distributed in the multidimensional space. The confidence intervals of the fitted parameters were estimated using lsqcurvefit in Matlab. In order to compare different kinetic models the standard deviation of the SSR was calculated by boostrap. Models that showed a SSR worse than 10 standard deviations with respect to the best model were discarded.

### Isothermal Titration Calorimetry

Isothermal titration calorimetry (ITC) assay was performed on a VP-ITC (Microcal VP-ITC (Malvern Panalytical) as previously described^7^. Briefly, titration was carried out at 15°C in a buffer containing 20 mM Tris-HCl pH 8.0, 0.3 M NaCl, 10 mM MgCl2 and 0.5 mM AMP-PNP. The concentration of dimeric DesKC was 15 μM and DesR_REC_ 380 μM. Raw data were analyzed with NITPIC v1.2.7^63,64^ and integrated binding isotherms were fitted to a model with two independent sequential sites using SEDPHAT v12.1b^65^. Plots including corrected thermograms, fittings of binding isotherms and residuals were done with GUSSI^66^.

## Data availability

The X ray structures presented have been deposited in the wwPDB with accession codes 7SSJ (DesK-DesR complex in the phosphatase state) and 7SSI (DesK-DesR_Q10A_ complex in the phosphotransfer state). Raw X ray diffraction data corresponding to each one of these structures are publicly available at SBGrid Data Bank (http://data.sbgrid.org) as dataset entries XXX.

## Acknowledgements

We thank Joaquin Dalla Rizza, Nicole Larrieux and Analia Lima for assistance in protein purification, crystallization and densitometric quantification, respectively. We acknowledge computational and storage services (Maestro cluster) provided by the Institut Pasteur IT Dept (Paris).

## Footnotes

1 REC: the phosphorylatable receiver domain of response regulators

