## Supplemental Figures and Tables for "Molecular basis of unidirectional information transmission in two-component systems: lessons from the DesK-DesR thermosensor"

**Extended Data Table S1**: X ray diffraction data collection and refinement statistics.

| | Phosphatase complex | $DesKC_{H188E}$:$DesR_{REC-Q10A}$ |
|---|---|---|
| **Data collection** | | |
| Space group | $P3_121$ | $P2_1$ |
| Cell dimensions | | |
| a, b, c (Å) | 94.69 94.69 240.85 | 88.07 115.63 91.37 |
| $\alpha$, $\beta$, $\gamma$ (°) | 90 90 120 | 90 116.7 90 |
| Resolution (Å) | 82.01-2.52 (2.73-2.52) | 34.41-3.41 (3.46-3.41) |
| Unique reflections | 34301 (1709) | 22399 (1046) |
| $R_{meas}$ | 0.085 (2.457) | 0.172 (1.599) |
| $R_{pim}$ | 0.020 (0.614) | 0.090 (0.858) |
| $CC_{½}$ | 0.999 (0.502) | 0.997 (0.446) |
| $I/\sigma I$ | 21.3 (1.3) | 7.5 (0.8) |
| Completeness (spherical) | 79.6 (18.9) | 99.6 (93.8) |
| Completeness (ellipsoidal) | 94.9 (66.9) | - |
| Redundancy | 17.8 (14.4) | 3.6 (3.3) |
| Refinement | | |
| Resolution (Å) | 33.89-2.52 (2.67-2.52) | 34.41-3.41 (3.49-3.41) |
| Number of refls used (N in the free set) | 1720 (38) | 20830 (24) |
| $R_{work}$ / $R_{free}$ | 0.253 / 0.289 | 0.252 / 0.283 |
| Number of atoms | | |
| Protein | 7955 | 8668 |
| Ligands + ions | 116 | 130 |
| Water | 22 | 10 |
| B factors (Å$^2$) | | |
| Wilson plot | 94.2 | 74.3 |
| Mean (overall) | 104.3 | 90.5 |
| R.m.s. deviations | | |
| Bond lengths (Å) | 0.008 | 0.008 |
| Bond angles (°) | 0.99 | 1.06 |
| Number of residues in Ramachandran plot[§] (favored/allowed/outliers) | 999/30/1 | 1048/37/1 |
| PDB ID | 7SSJ | 7SSI |

**Extended Data Table S2.** Reaction coordinate definitions and initial values for all the MSMD simulated reactions.

| **Phosphotransfer reaction** | **d (His N - P)** | **d (P - Asp O)** | **RC (initial)** | **RC (final)** |
|---|---|---|---|---|
| Desk-DesR (Phospo His188 - Asp54) | 1.75 | 3.85 | -2.15 | 2.15 |
| YPD1-SLN1/R1 (Phospho His64 - Asp1144) | 1.75 | 3.05 | -1.3 | 1.3 |
| solution (Phospho His - Asp ) | 1.75 | 5.25 | -3.5 | 3.5 |
| **Phosphatase reaction** | **d (Wat O - P)** | **d (P - Asp O)** | **RC (initial)** | **RC (final)** |
| DesK-DesR (Phospho Asp54) | 3.75 | 1.6 | -2.15 | 2.15 |
| DesR auto-phosphatase (Phoshpo Asp54) | 3.75 | 1.6 | -2.15 | 2.15 |

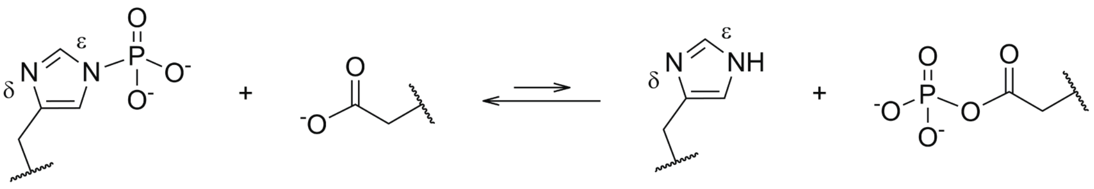

**Extended Data Figure 1**: Schematic illustration of the phosphoryl-transfer reaction.

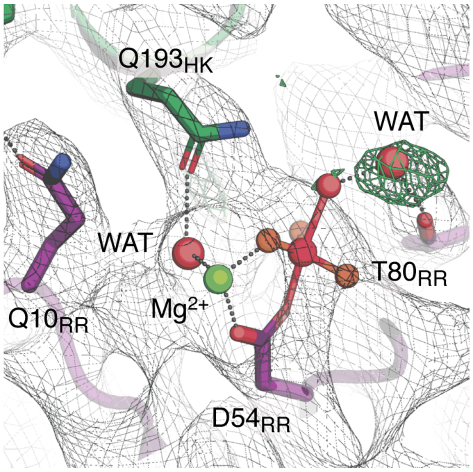

**Extended Data Figure 2**: Modeling of the $MgF_3^-$ transition state in the reaction center of the DesK-DesR complex in the phosphatase state. 2mFobs-DFcalc electron density map contoured at 1σ is shown as a gray mesh and in green positive peaks of the mFobs-DFcalc electron density contour at 3.5σ. Key residues are depicted in sticks.

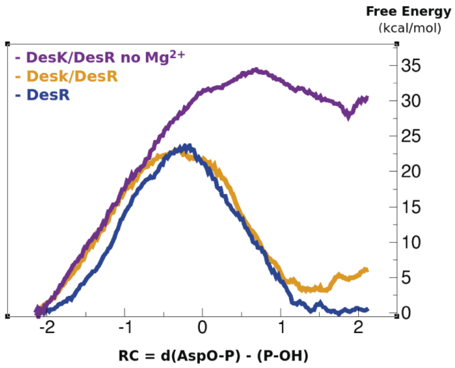

**Extended Data Figure 3**: Energy profile of the DesK phosphatase-catalyzed reaction in the presence (orange) and absence of $Mg^{2+}$ (magenta). The blue line shows calculations corresponding to the dephosphorylation of DesR in the absence of DesK.

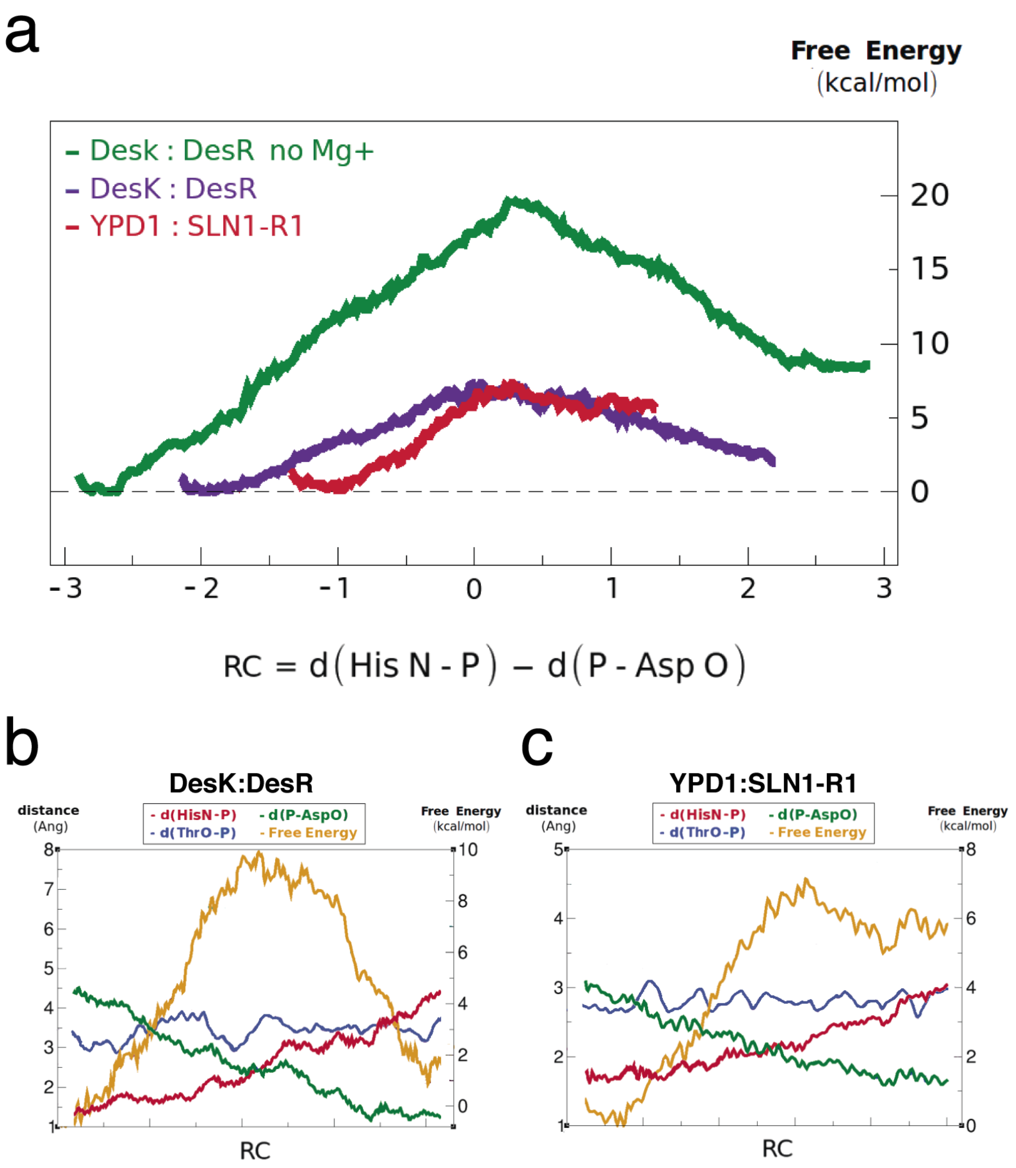

**Extended Data Figure 4:** Phosphoryl-transfer reaction QM/MM steered molecular dynamics calculations. **a)** Free Energy profile as a function of the reaction coordinate for the DesK-DesR system in the presence (purple) or absence of $Mg^{2+}$ (green). The red line corresponds to Ypd1:Sln1-R1. **b** and **c)** Main interatomic distances fluctuations as a function of the reaction coordinate for the DesK:DesR system (**b**) and Ypd1:Sln1-R1 (**c**) systems.

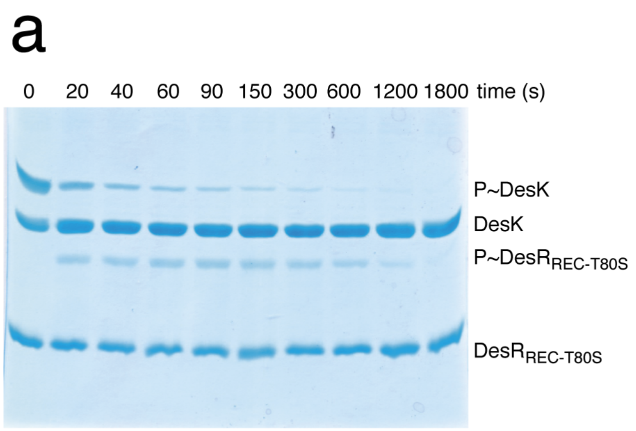

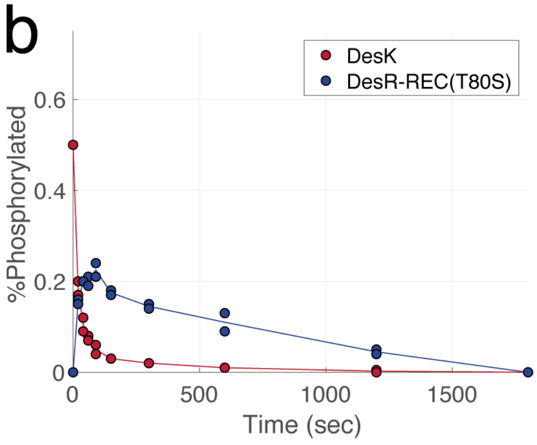

**Extended Data Figure 5**: a) Representative PhosTag SDS gels analyzed for the DesKC~P and $DesR_{REC\text{-}T80S}$ phosphoryl-transfer assay. B) Densitometry analysis of the phosphoryl-transfer assay between DesKC~P and $DesR_{REC\text{-}T80S}$.

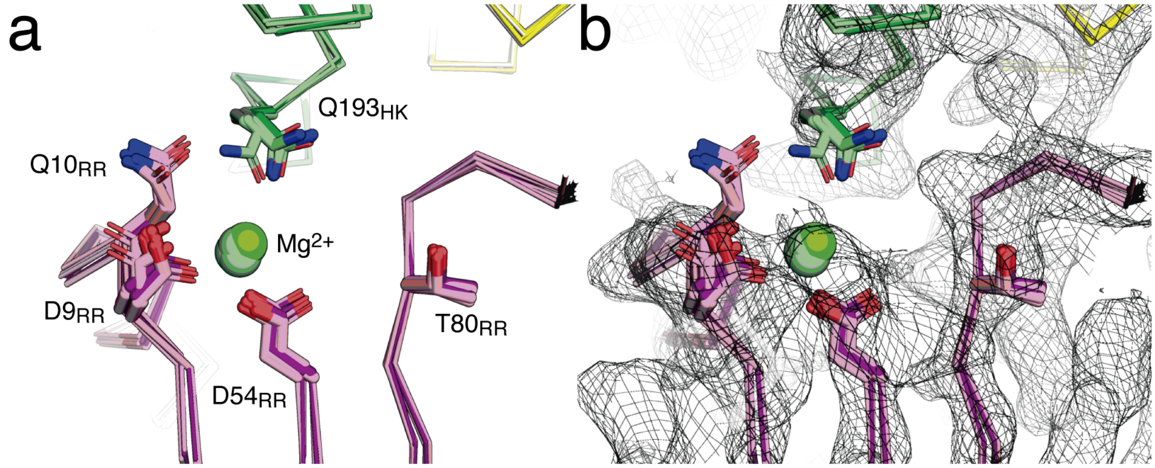

**Extended Data Figure 6**: Modulation of phosphoryl-transfer reversibility. (a) $DesK_{H188E}$:$DesR_{REC\text{-}Q10A}$ crystal structure (the two independently refined complexes of the asymmetric unit are depicted in purple) was aligned to $DesK_{H188E}$:$DesR_{REC}$ structure (colored as light magenta, PDBId 5IUK) using residues 190 to 230 of the DHp domain of DesK (in green and yellow) as a reference. (b) Similar view as in (a), showing the 2mFobs-DFcalc electron density contour at 1σ of the $DesK_{H188E}$:$DesR_{REC\text{-}Q10A}$ crystal structure.

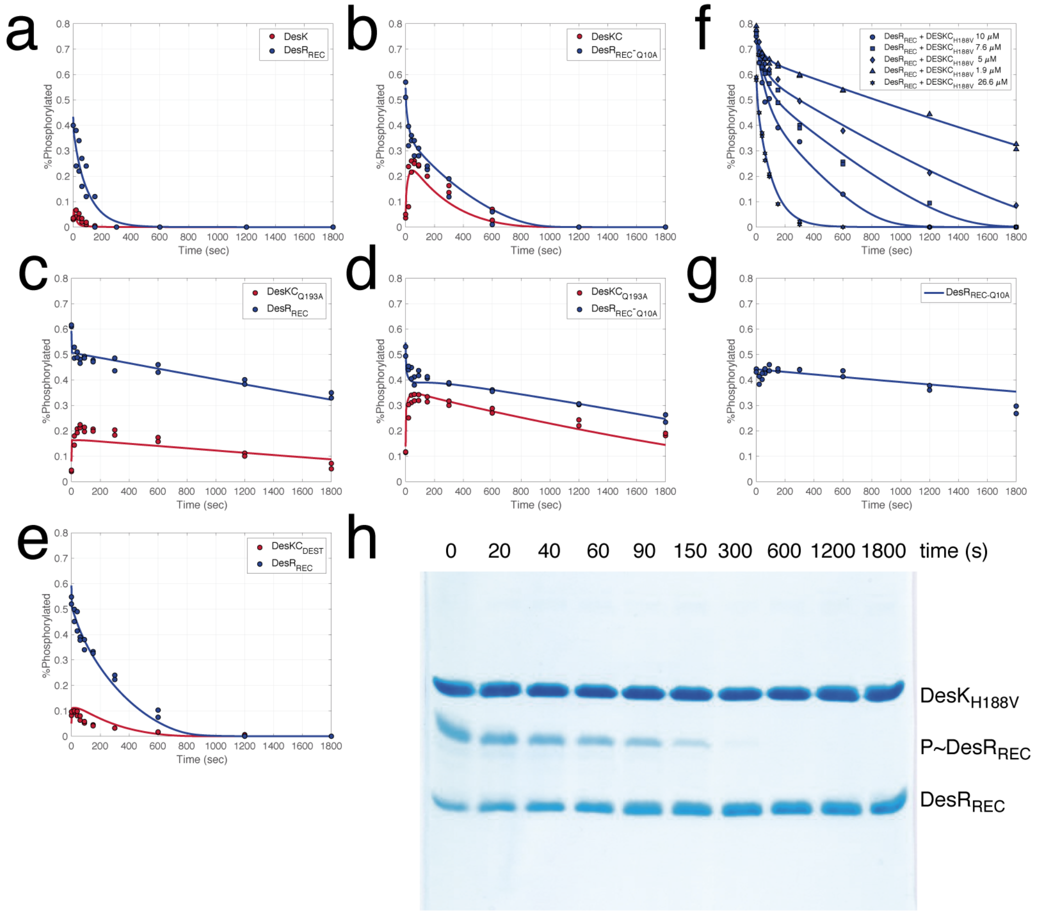

**Extended Data Figure 7**: Dephosphorylation assay starting the reaction either with P~$DesR_{REC}$ (a, c, e and f) or P~$DesR_{REC\text{-}Q10A}$ (b, d and g) in the presence of DesKC (a, b), $DesKC_{Q193A}$ (c, d), $DesKC_{DEST}$ (e) or $DesKC_{H188V}$ (f). The continuous blue or red traces represent the simulations from the best-fitted model. h) Representative PhosTag SDS gels for the phosphatase assay, starting with P~$DesR_{REC}$ (26 $\mu$M) and $DesKC_{H188V}$ (26 $\mu$M).

a

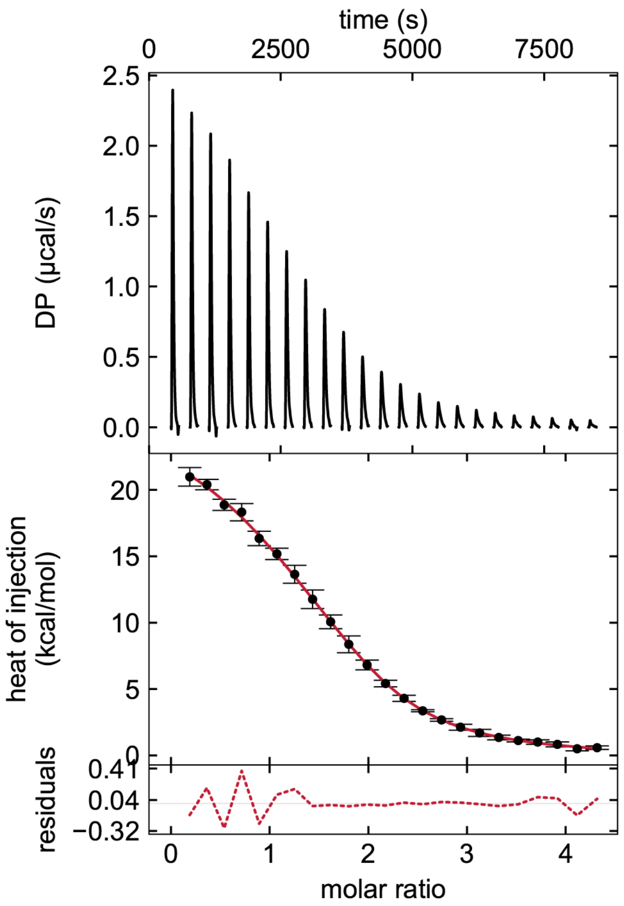

b

| | Ka ($M^{-1}$) | ΔG (kcal.mol$^{-1}$) | ΔH1 (kcal.mol$^{-1}$) | T.ΔS ($x10^3$ kcal.mol$^{-1}$) |
|---|---|---|---|---|
| DesKC | $1.21x10^6$ | -8.02 | 25.53 | 33.55 |
| | $3.15x10^5$ | -7.24 | 8.63 | 15.87 |

**Extended Data Figure 8**: Isothermal titration calorimetry (ITC) of DesKC:$DesR_{REC}$. a) Top panel shows the raw heat flow data over time of the isothermal titration assay; bottom panel shows integrated heat exchange as a function of $DesR_{REC}$ (monomer): DesKC (dimer) molar ratio. b) Isothermal titration calorimetry parameters.

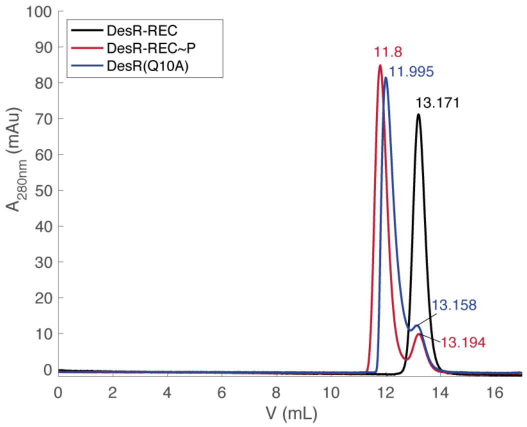

**Extended Data Figure 9**: Size exclusion-chromatography comparing of P~DesR$_{REC}$ (red), DesR$_{REC}$ (black) and P~DesR$_{REC-Q10A}$ (blue). Phosphorylation of DesR$_{REC}$ and DesR$_{REC-Q10A}$ was performed by incubating the proteins with acetyl-phosphate for 1 hour in the presence of 20 mM $MgCl_2$ at room temperature. The degree of phosphorylation of both proteins was calculated to be of 60%, as evaluated by PhosTag SDS gels.

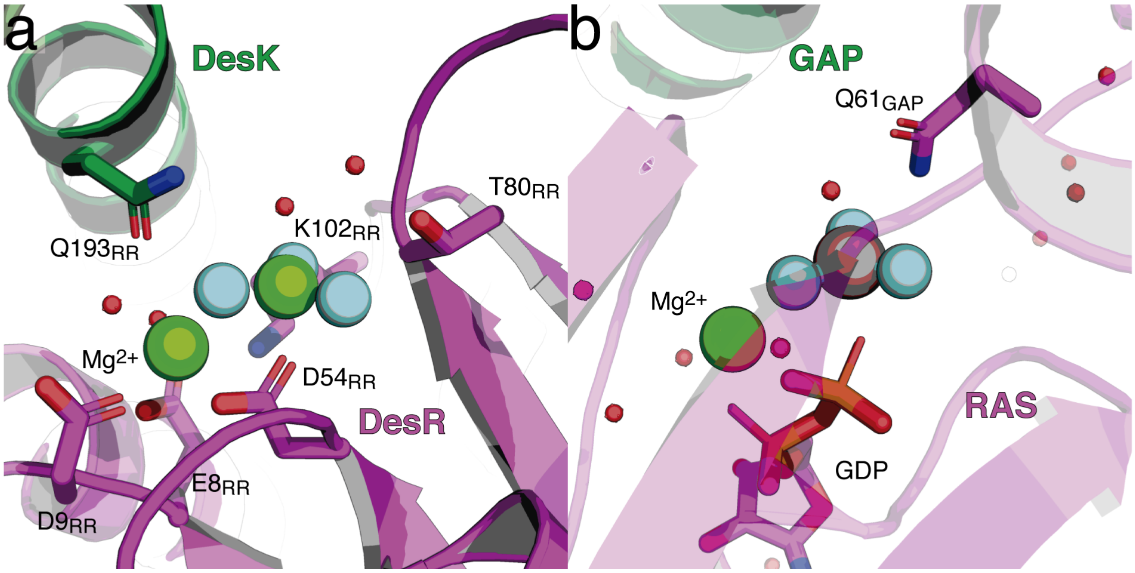

**Extended Data Figure 10**: Comparison of the reaction centers of the phosphatase state of the DesK:DesR (a) and the Ras:Gap (b, PDBid 1WQ1) complex. a) DesK is depicted in green showing the key residue Gln193 in sticks. DesR is colored as magenta and the $Mg^{2+}$ and $MgF_3^-$ are shown as spheres. Highly conserved residues Asp54, Glu8 and Asp9 are displayed as sticks and water molecules as small red spheres. b) Similarly to panel a), Gap is colored in green and Ras in magenta. Gln61 and GDP are displayed as stick and in sphere are shown the transition state mimetic $AlF_3^-$ and the $Mg^{2+}$.

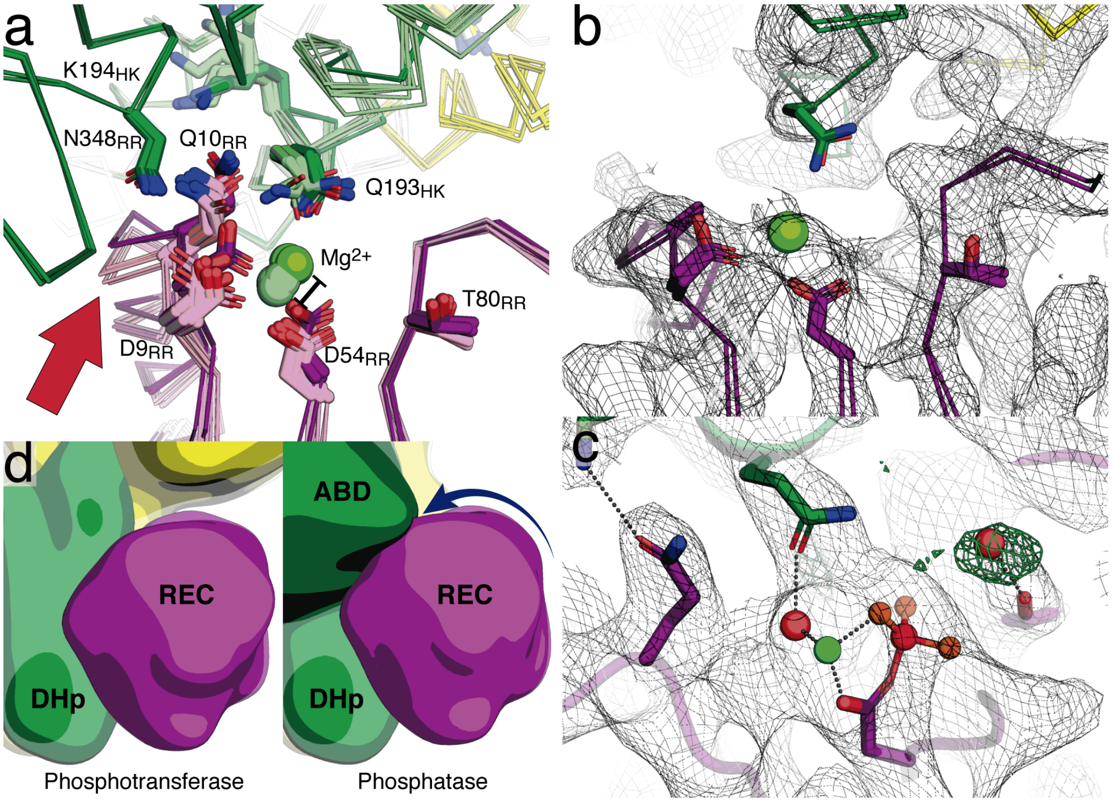

**Extended Data Figure 11**: Futile cycle minimization through guided repositioning of the RR. a) Structural alignment of phosphatase vs phosphotransferase complexes using residues 190 to 230 of the DHp of DesK as a reference. The two chains of the dimer of DesK are shown in green and yellow and DesR is colored as magenta. Phosphatase state is displayed in darker color and phosphotransferase structure is shown in light tones. The red arrow indicates the direction of the small shift comparing the structures. b) and c) Similar view as panel a) showing the electron density maps contoured at 1σ of the phosphotransferase (PDB id: 5IUK) (b) and phosphatase (PDB id: 7SSJ) (c) complexes. d) Schematic representation of the shift experienced by the RR through interaction with the ABD.

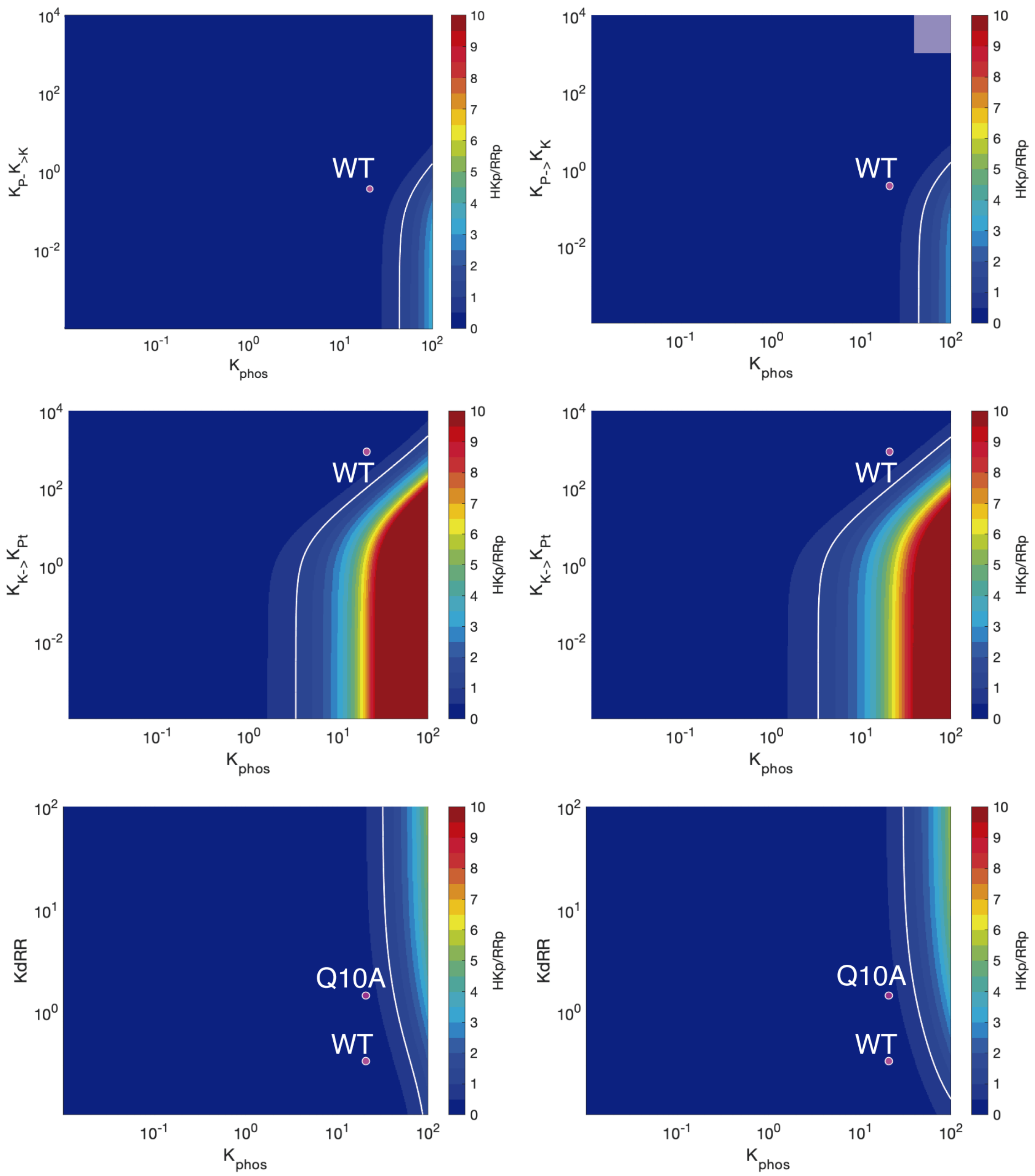

**Extended Data Figure 12**: Analysis of key unidirectional determinants. Phosphorylation distribution between the HK and RR in the absence of phosphatase activity (equivalent to using $DesK_{Q193A}$ in the phosphotransfer assays). The first column shows the analysis starting the simulation with the phosphorylated HK, whereas the second column displays reaction starting from the phosphoryl-moiety attached to the RR. The simulation is run for 600 seconds and the ratio of the [HK~P]/[RR~P] is calculated for different combination of parameters. As a reference we show the experimental determination using wild type and $DesR_{Q10A}$ mutant (magenta dots).

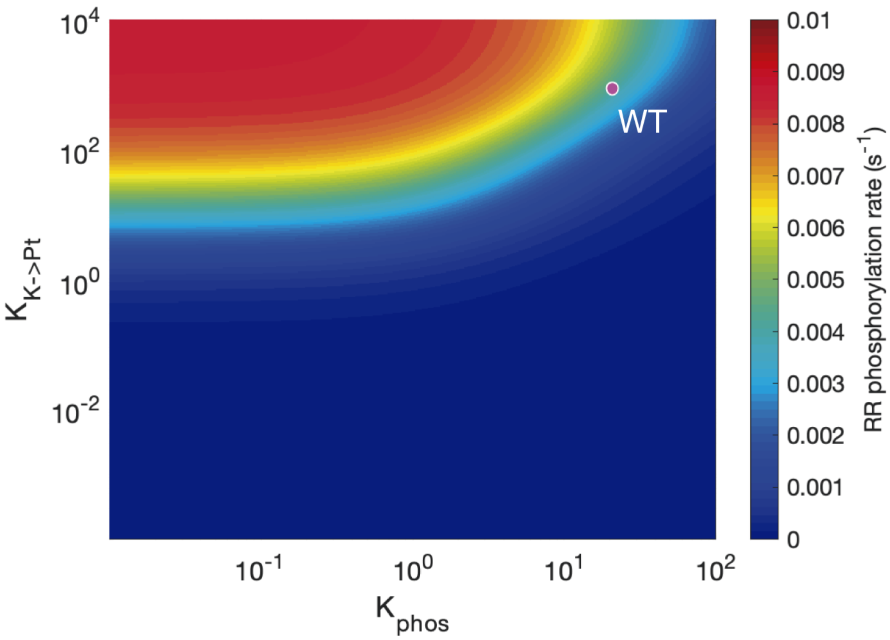

**Extended Data Figure 13**: HK catalyzed RR phosphorylation efficiency. In the presence of the signal ($K_{P\text{-}K}$ << 1) the rate of RR phosphorylation was simulated using kinase autophosphorylation kinetic constants[29] and using different combinations of parameters describing the reversibility of phoshoryl-transfer (Kphos = k4/k3) and the kinase/phosphotransferase conformational equilibrium ($K_{K\text{-}Pt}$ = *k8b*/*k7b*). Magenta dot corresponds to the best fitted parameters for the wild type DesKC and $DesR_{REC}$.

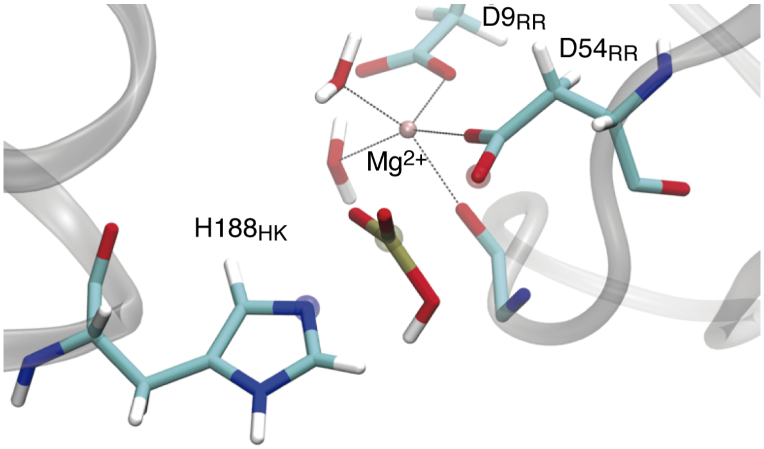

**Extended Data Figure 14**: QM region for phosphotransfer reaction QM-MM simulations.
